# Early ingestive experience with a high-fat diet tunes satiation and nutrient-specific appetitive behaviors

**DOI:** 10.64898/2026.02.24.707784

**Authors:** Meaghan McCoy, Ciorana Roman-Ortiz, Jason Perez, Lindsey A. Schier, Anna K. Kamitakahara

**Author notes:** **Correspondence:** Anna Kamitakahara, PhD, Children’s Hospital Los Angeles, 4661 Sunset Blvd, MS#135, Los Angeles, CA 90027. These authors contributed equally to this work (Meaghan McCoy, Ciorana Roman-Ortiz).

## Abstract

Overconsumption of foods rich in fats, sugars, and calories is a major contributing factor to increased risk for cardiometabolic disease. Ingestive experience with these foods can begin early in children, yet there is limited understanding of the impact of early life nutrition on the development of vagal afferent neurons necessary for coordinating appetitive and satiating behaviors. To address this gap, mice reared on a chow diet (control) were compared to those reared on a high-fat diet (HFD^EARLY^). We demonstrate that the vagally-mediated satiation response to cholecystokinin (CCK) does not mature until adolescence in chow-reared mice. However, HFD^EARLY^ exposure triggers a precocious maturation of this response, accompanied by transcriptomic changes in the nodose ganglion. Durable changes in appetitive behaviors were also evident in adult HFD^EARLY^ mice, which consumed more lipid than control mice. Behavioral analyses point to alterations in orosensory processing and enhanced appetition in adult HFD^EARLY^ mice. Furthermore, female HFD^EARLY^ mice showed a selective increase in consumption of a calorie-dense liquid diet in adulthood, accompanied by changes in body composition, suggesting a lasting, sex-specific vulnerability to overconsumption of palatable, energy-dense foods. Together, these results establish nutrient exposure as a significant contributor to vagal circuit maturation and function.

## Introduction

Early life represents a period of significant learning about food, including how much to consume based on the internal sensation of fullness and which foods are favored. Vagal sensory neurons are a key component of the neural circuits coordinating the opposing ingestive drives of appetition and satiation, which stimulate and inhibit consumption, respectively. Nutrient-responsive vagal sensory afferents in the gastrointestinal tract make multi-synaptic connections to ascending dopaminergic circuits to generate appetitive responses that reinforce consumption of fats and sugars through integration with orosensory cues (e.g. taste, mouthfeel)^1–9^. Vagal afferents also convey nutrient-driven and mechano-responsive satiation signals through ascending hypothalamic circuits to coordinate meal termination^5,10,11^. While signaling of vagal-dependent appetition and satiation have been well delineated in the adult nervous system, limited information is available on when and how these behaviors develop and mature.

In rodents and humans, vagal sensorimotor circuits undergo a prolonged period of growth and maturation that begins prenatally and extends into postnatal life as ingestive experience is gained^12–17^. In rodents, vagal sensory neurons are born by day 9 of gestation and reach targets in the stomach and small intestine by embryonic day 13.5 and 16.5, respectively^12,18^. Ascending vagal projections to central targets in the nucleus of the solitary tract (NTS) are also present at birth and undergo continued synaptogenesis through postnatal day (P) 10^13,18^. Despite reaching central and peripheral targets prenatally, substantial differences in gene expression are also observed after P7, particularly in genes related to axon guidance and synaptic potentiation, highlighting the possibility that vagal sensory circuits continue to mature past the first postnatal week^14^. Notably, expression of the cholecystokinin (CCK) receptor (*Cckar*) increases dramatically following birth, increasing nearly 3-fold after P7 in mice^14^.

Vagal nutrient sensing pathways critically depend on gut-derived CCK for signaling satiation and appetitive reward^1–3,5,7,10,11,17,19,20^, suggesting that these behaviors likely remain immature in early postnatal life. This is consistent with data in rats suggesting that nutrient sensation-based satiation signaling begins to arise during the third postnatal week^21^. As vagal sensory neurons grow and mature, they may be particularly vulnerable to perturbation by the dietary environment. Yet, there are critical knowledge gaps in understanding how the sensitivity of responses to vagally-mediated satiation are established developmentally, and whether early dietary exposure to energy-dense foods alters the formation of mature vagal responses in ways that promote overconsumption.

To address this gap, we used a well-established model of early life high-fat diet (HFD) exposure^22–25^ in which dams were provided with *ad libitum* access to a standard chow diet or to a high-fat diet from the birth of their litter until weaning, when all pups were then provided the chow diet for the remainder of the study. We tracked the impact of early life HFD on the developmental trajectory of behavioral sensitivity to the satiation hormone CCK and identified accompanying changes in the nodose transcriptome. A series of behavioral tests was then used to examine nutrient preference, appetition, and taste motivation in adulthood. Our results reveal that early dietary experience accelerates the functional maturation of vagal satiation circuits, produces lasting sex-specific alterations in appetitive behavior and susceptibility to overconsumption of calorie-dense foods, and establishes vagal circuit maturation as a target of early nutritional programming with lasting consequences for metabolic health.

## Results

### Timeline of Maturation for CCK-dependent Satiation and Appetitive Responses to Glucose

Cholecystokinin (CCK) is released from the intestinal epithelium in response to ingested nutrients and signals satiation (meal termination) through activation of vagal sensory neurons (VSNs) that project from the gut to the hindbrain^11,20,26–28^. CCK receptor expression within VSNs increases substantially during postnatal development, suggesting that CCK-dependent behaviors mature postnatally^14^. To directly test the maturation of satiation responses to CCK, male and female mice were longitudinally tested at several ages − postnatal days (P)35, P45, P60, and P120. At each age mice were fasted overnight, then given an injection of either saline or CCK the following morning along with pellets of a highly palatable high-fat diet to consume. At P45, P60, and P120, CCK injection elicited strong, highly consistent decreases in food intake (Figure 1A and B). By contrast, at P35 mice exhibited highly heterogenous satiation responses with some mice consuming very little food while others seemed largely unaffected by CCK injection. Using P120 as the mature response, all other ages were compared by one-way ANOVA with Dunnet’s correction for multiple comparisons. A statistically significant difference in the response at P35 was identified, but not at older ages, suggesting that mature responses to CCK injection develop by P45 (mid-late adolescence).

**Figure 1.**
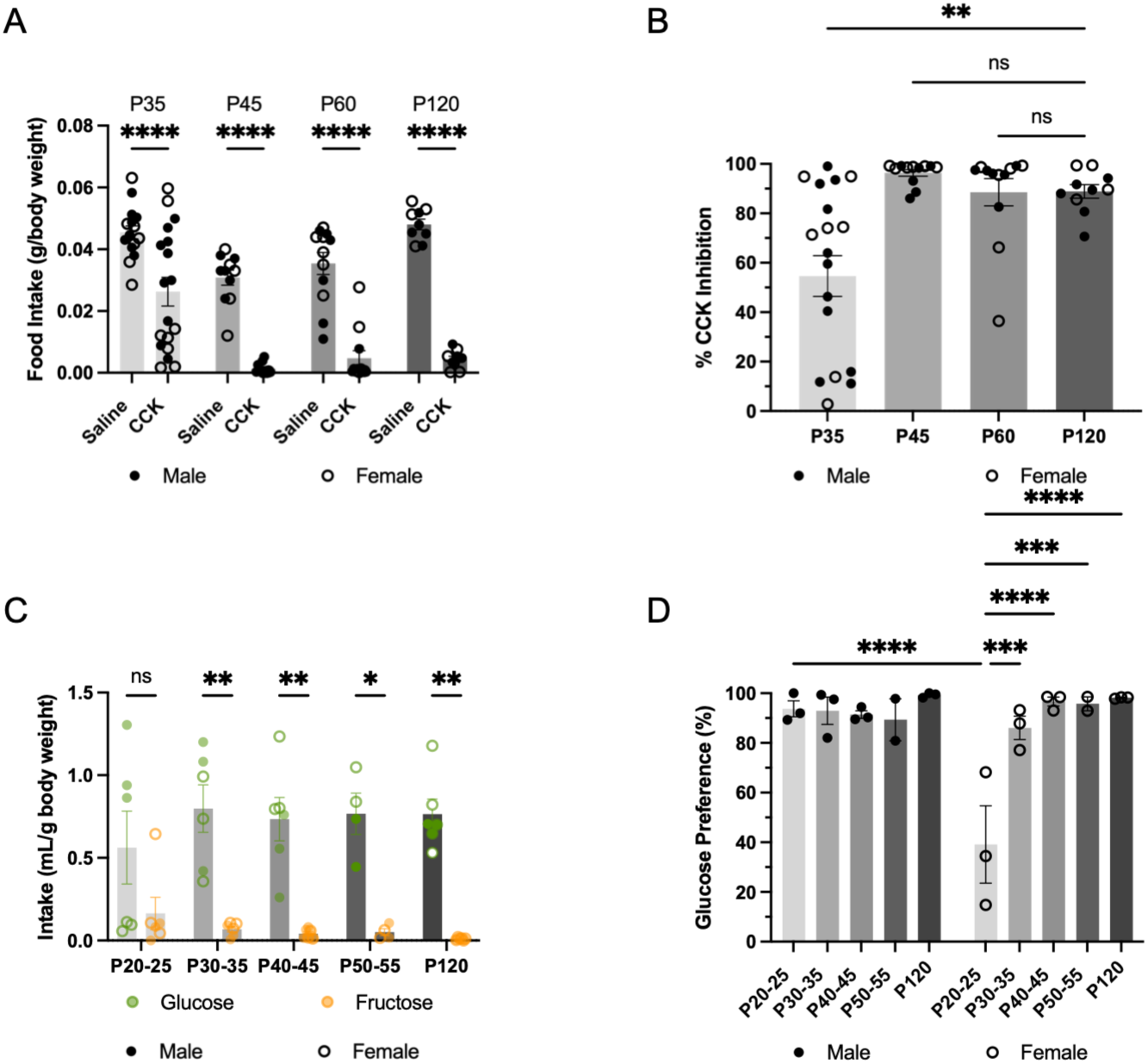
Maturation of CCK-dependent satiation and post-ingestive appetitive responding to glucose. A) HFD intake measured in male and female mice following injection with saline or CCK (10 ug/kg) at P35, P45, P60, or P120. Of note, body weight at P35 and P45 did not substantially differ, and were thus injected with the exact same dose and volume. A one-way ANOVA with Dunnet correction for multiple comparisons was used to analyze differences in food intake between each age compared to P120 (n=8-12). B) Percent CCK feeding inhibition across ages in male and female mice. A one-way ANOVA with Dunnet correction was used to analyze differences in percent inhibition between each age compared to P120. C) Intake of glucose and fructose solutions during the two-bottle preference test on the final test day across developmental ages (P20-25, P30-35, P40-45, P50-55, P120) in male and female mice (n=5-6 per age). Solutions were calorie-matched (0.34 kcal/mL). Amount consumed is normalized to the combined total body weight of the mice in each cage. D) Glucose preference (glucose intake as a percentage of total fluid intake) across the same developmental ages. A one-way ANOVA with Tukey correction was used to compare each age to P120 as the mature response. **** p<0.0001; ** p<0.01; * p<0.05; ns, not significant.

VSNs are also critical for driving post-ingestive appetitive responses^1,2,6,7,29^. To probe the maturation of post-ingestive appetitive behavior, a two-bottle preference test was administered between P20-25, P30-35, P40-45, P50-55, and around P120. At each age mice were pre-exposed to solutions of 10% glucose and 10% fructose delivered one at a time. Then on the test day the mice were provided both solutions to assess preference. Solutions in each bottle were matched for caloric content and sweet taste. In mature mice, glucose is known to positively reinforce intake from a postingestive site of action, while fructose does not^7,29–31^. We replicated this behavior demonstrating that there is a robust preference for glucose by P35 in both sexes that is retained in adult mice (Figure 1C and D). However, when tested between P20-25, a heterogeneous glucose preference response was observed, driven by reduced preference for glucose specifically in females. Using P120 as the mature response, all other ages were compared by one-way ANOVA with Dunnet’s correction for multiple comparisons. A statistically significant difference in the glucose preference response in females between P20-25 was identified, but not at older ages, suggesting that mature appetitive responses to sugars develop by P25 in males but not until P35 in females.

### Early Life Dietary Manipulation

Given this protracted maturational timeline, we next sought to test the effect of dietary experience on the maturation of CCK-dependent satiation signaling. Prior studies have demonstrated that HFD consumption in adulthood results in impaired satiation responses to CCK^32,33^. To examine whether satiation responses are similarly impaired by early life ingestive experience, we used a model of HFD exposure limited to the lactation and complimentary feeding period. Dams were randomized into groups receiving either a control chow diet or a HFD (HFD^EARLY^) for 3 weeks beginning on the day of birth of the litter until weaning (Supplemental Figure 1A). HFD consumption during the lactation period is known to increase lipid content of the milk and weight gain in offspring^34,35^. Consistent with prior studies, male and female HFD^EARLY^ pups exhibited a significant increase in body weight compared to sex-matched controls by P21 (Supplemental Figure 1B and E). At P21, all mice were weaned onto a standard chow diet. By P35, two weeks after weaning onto chow, body weight did not differ between control and HFD^EARLY^ mice. However, a 20% increase in adiposity remained in both male and female HFD^EARLY^ mice compared to sex and age matched control mice (Supplemental Figure 1C and F). The percent lean mass was also significantly decreased in HFD^EARLY^ females, but not males at P35 (Supplemental Figure 1D and G).

### Impact of Early Life Diet on the Development of CCK-dependent Satiation Responses

The impact of early dietary experience on CCK-mediated satiation signaling was examined at P35 in mice exposed to HFD in early life and control, chow-reared mice. Following saline injection, no statistically significant differences in food intake were detected between groups (Figure 2B and F). As expected, CCK injection resulted in a reduction in food intake in control mice that was highly heterogeneous. By comparison, HFD^EARLY^ mice exhibited robust, consistent decreases in food intake in response to CCK at this age. Both male and female HFD^EARLY^ mice consumed significantly less food than sex-matched control mice (Figure 2B and F) and had a greater percent inhibition when assessing each individual animals’ differential response to saline vs CCK (Figure 2C and G). The data suggest that early life HFD exposure promotes precocious maturation of CCK-dependent satiation responses.

**Figure 2.**
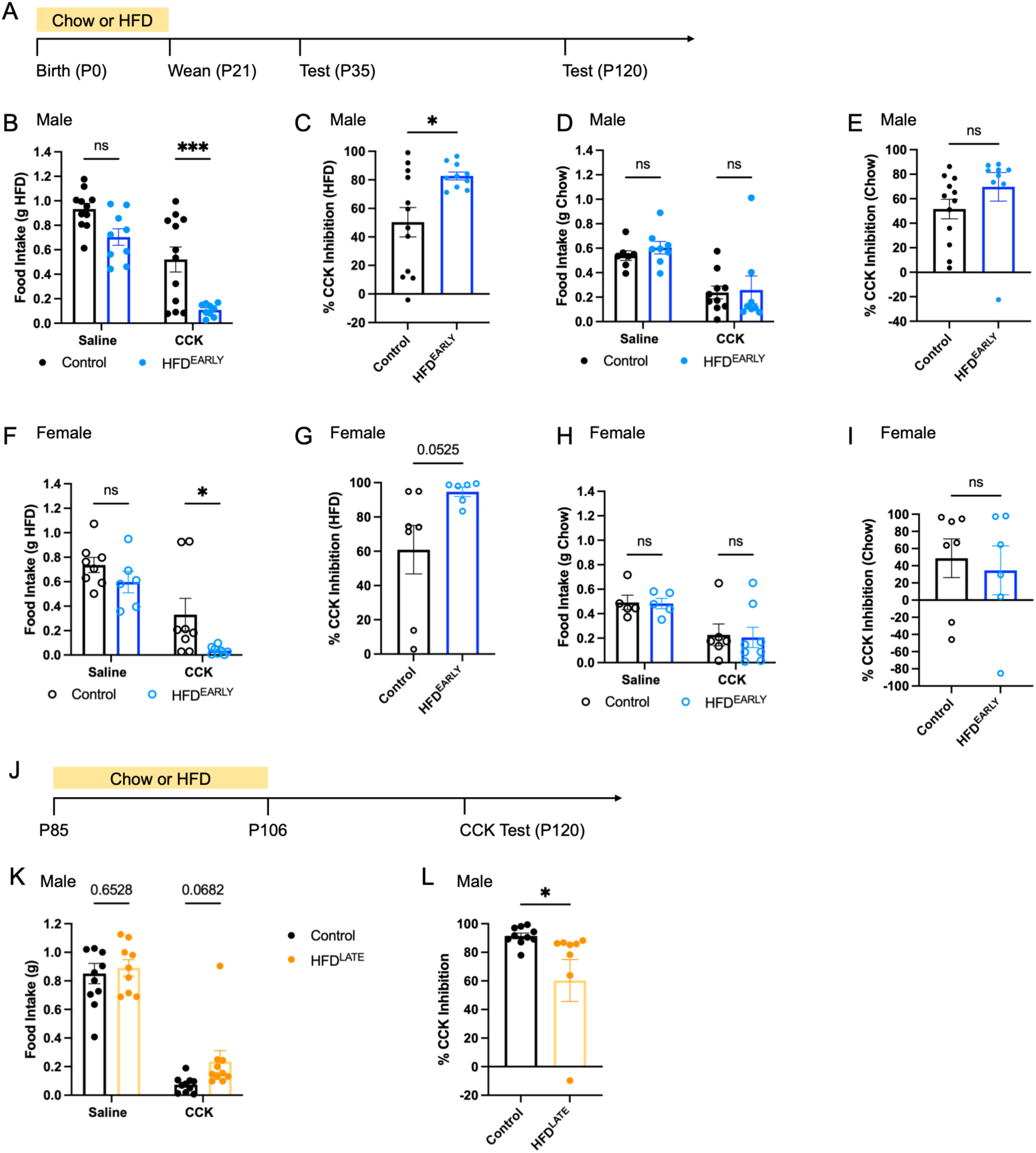
HFD feeding in early life drives precocious maturation of CCK-mediated satiation responses. A) Timeline for the HFD^EARLY^ dietary manipulation and testing. B) Food (HFD) intake in males at P35, 30 minutes following saline or CCK injection. C) Percent feeding inhibition calculated from each individual male animal’s saline and CCK response tested with HFD. D) Food (chow) intake in males at P35 following saline or CCK injection. E) Percent feeding inhibition calculated from each individual male animal’s saline and CCK response tested with chow. F) Food (HFD) intake in females at P35, 30 minutes following saline or CCK injection. G) Percent feeding inhibition calculated from each individual female animal’s saline and CCK response tested with HFD. H) Food (chow) intake in females at P35 following saline or CCK injection. I) Percent feeding inhibition calculated from each individual female animal’s saline and CCK response tested with chow. J) Timeline for the HFD^LATE^ dietary manipulation and testing. K) Food (HFD) intake in adult males, 30 minutes following saline or CCK injection. L) Percent feeding inhibition calculated from each individual male animal’s saline and CCK response. One extreme value was observed in Figure 2L; repeating the analysis with and without this data point did not alter the conclusion (p=0.022 with inclusion; p=0.048 with exclusion). A two-way mixed-effects ANOVA was used to analyze differences in response to saline or CCK between groups. A two-tailed unpaired t-test was used to analyze differences in percent feeding inhibition between groups (* p<0.05, *** p<0.001). ns, not significant.

In addition to testing with highly palatable HFD pellets, the same CCK injection paradigm was used to examine the satiation response to standard chow consumption at P35. Following CCK injection, chow intake was similar comparing control and HFD^EARLY^ mice, although a trend toward increased inhibition was observed in male but not female HFD^EARLY^ mice (Figure 2D, E, H, and I). The combined results of testing using chow and HFD suggest that the nutritional content of the test diet moderates the feeding response in HFD^EARLY^ mice.

Results obtained following early life HFD exposure run counter to the results of prior studies completed in adult rodents in which CCK-dependent satiation signaling is impaired by HFD exposure^32,33^. To directly compare phenotypes observed in early vs late models of HFD exposure, a small replication cohort of adult male mice was assessed. Adult mice were provided with either a control chow diet or a HFD for 3 weeks (HFD^LATE^), then switched to chow for 2 weeks to align with the timing of the dietary manipulation applied in early life (Figure 2J). Satiation responses to CCK were tested in control and HFD^LATE^ mice provided with HFD. Following saline injection, both groups consumed a similar amount (Figure 2K). Following injection with CCK, a trend toward increased food intake was observed in HFD^LATE^ mice compared to controls, along with a significant decrease in percent feeding inhibition (Figure 2K and L). While one extreme value was observed among CCK injected HFD^LATE^ mice, repeating the analysis in Figure 2L with and without this data point did not alter the conclusion (p = 0.022 with inclusion; p = 0.048 with exclusion). This HFD^LATE^ cohort demonstrates that the effect of HFD exposure in early life differs from the response to HFD in the mature nervous system.

To examine the mechanisms through which early life diet alters responses to CCK, we performed highly multiplexed *in situ* hybridization (HiPlex RNAscope) to quantify CCK receptor (*Cckar*) expression, as well as expression of other GI-projecting cell type specific markers. Previous studies completed in adult mice fed a HFD suggest that impaired responsiveness is mediated via reductions in CCK receptor expression in the nodose ganglion^32^. Therefore, we hypothesized that *Cckar* expression would be altered in HFD^EARLY^ mice at P35, given their precocious maturation of satiation responses to CCK. However, no difference was detected in the proportion of cells expressing *Cckar*, nor the mean expression of *Cckar* per cell (Supplemental Figure S2). Additionally, no difference was observed in *Cckar* expression within subsets of neurons expressing any specific cell type marker (*Trpa1*, *Vip*, *Calca*, *Sst*, *Oxtr*, *Cartpt*, *Glp1r*, *Npy2r*, *Lepr*, *Gpr65*, or *Htr3b*). The lack of differential expression of *Cckar* between groups suggests that the mechanisms through which CCK-dependent satiation signaling are modulated following HFD consumption differ in early life from the mechanisms employed in the mature nervous system.

To identify novel candidate molecular mechanisms that mediate the precocious CCK-dependent satiation response observed following early life HFD, bulk RNA sequencing was performed using nodose ganglion samples from mice at P21 and P35 following rearing on chow or HFD. In addition, these samples were analyzed together with previously sequenced nodose ganglion samples from P7, P50, and P90 chow reared mice to more broadly understand the developmental trajectory of vagal sensory neuron development^14^. Principal component analysis revealed that samples clustered by age following a clear temporal trajectory (Figure 3A, indicated by arrows). Of the samples collected from control mice, P7, P21, and P90 were the furthest distance apart in principal components space, suggesting that these ages are the most transcriptionally distinct from one another. Further differential analysis of gene expression between these ages provided context for developmental processes occurring between each age, such as expression of genes related to cell proliferation, cell death, myelination, and synaptic vesicle production, among others (Supplemental Figure S3, see methods for greater detail). The expression of genes related to cell cycle/proliferation, cell death/apoptosis, myelin production, and extracellular matrix production were all most highly expressed at P7 compared to later ages. By contrast, expression of genes related to cytoskeletal organization, post-synaptic density proteins and ligand gated ion channels, G proteins, and cytokine receptors peaked at P21 before decreasing in adult samples, suggesting that the time around weaning is critical for expression of these groups of genes largely involved in being responsive to afferent signals. Gene expression related to ion pumps and ion channels peaked by P21, then remained constant suggesting that membrane polarization properties likely stabilize around weaning. Lastly, expression of genes related to synaptic vesicle production increased significantly by P21, but continued to increase modestly by P90, suggesting that maturation of efferent signaling continues after P21, consistent with the timeline for maturation of CCK-dependent satiation demonstrated in Figure 1. Together, this analysis of gene expression over time provided a foundation for understanding subsequent differential expression of genes between control and HFD^EARLY^ nodose ganglion samples.

**Figure 3.**
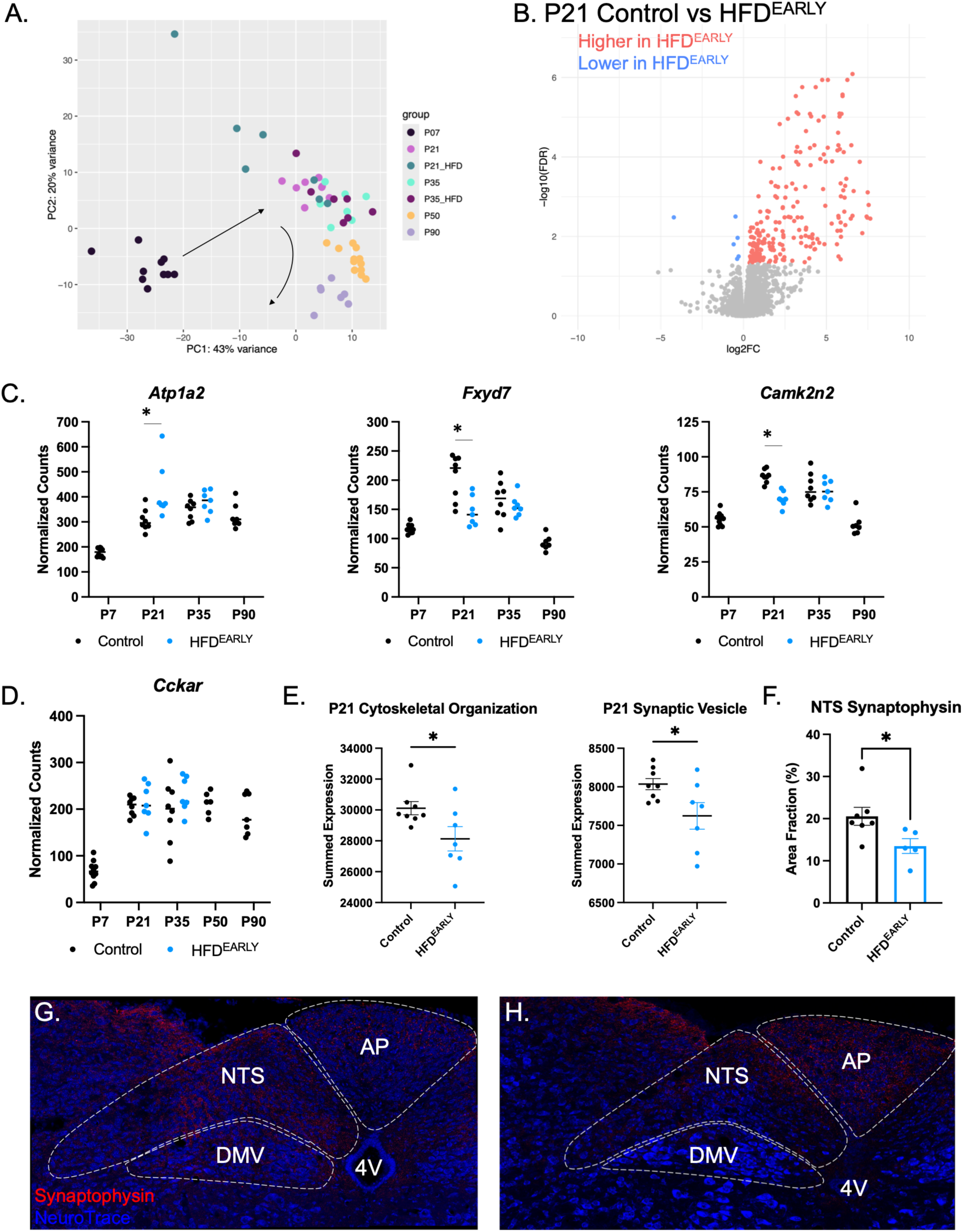
Early life diet alters gene expression in the developing nodose ganglion and synaptic structure in the NTS. A) Principal component analysis (PCA) of sequenced nodose ganglion samples from control chow-reared mice at P7, P21, P35, P50, and P90, compared to HFD^EARLY^ mice at P21 and P35. Arrows indicate a clear developmental trajectory of gene expression through PCA space. B) Volcano plot of differentially expressed genes (DEGs) in HFD^EARLY^ mice compared to control mice at P21. Genes higher in HFD^EARLY^ mice are shown in red; genes lower in HFD^EARLY^ mice are shown in blue. C) Normalized counts for three genes modulating neuronal excitability − *Atp1a2*, *Fxyd7*, and *Camk2n2* − that were differentially expressed between control and HFD^EARLY^ mice at P21 (adjusted p values calculated in DESeq2). D) Normalized counts for *Cckar* (CCK receptor) across ages and dietary groups, demonstrating that *Cckar* expression does not differ between control and HFD^EARLY^ mice at any age examined (n=10 P7; 7-8 P21; 7-8 P35; 6 P50; 7 P90). E) Summed expression of genes in the cytoskeletal organization and synaptic vesicle domains at P21 in control and HFD^EARLY^ mice. F) Area fraction of synaptophysin-immunoreactive puncta in the nucleus of the solitary tract (NTS) at P21 in control and HFD^EARLY^ mice. A two-tailed unpaired t-test was used for comparisons in panels E and F. * p<0.05. G-H) Representative images of immunolabeling for synaptophysin (red) with NeuroTrace (blue) in the NTS of P21 control (G) and HFD^EARLY^ (H) mice.

Comparison of control and HFD^EARLY^ nodose ganglion samples at P21 identified 176 differentially expressed genes (DEGs), as well as distinct clustering of P21 HFD^EARLY^ samples in principal components space (Figure 3A and B). By comparison, only 4 DEGs were identified between HFD^EARLY^ and control nodose ganglion samples at P35, suggesting that despite the differential responsiveness to CCK observed at this age that many of the transcriptional changes that mediate this response may have occurred earlier, likely during the diet manipulation period. Therefore, we focused further analyses on the DEGs identified at P21. Graphic representation of the DEGs using a volcano plot demonstrates that the vast majority of DEGs were higher in HFD^EARLY^ mice compared to chow reared controls (Figure 3B). Consistent with the HiPlex results described above, no difference in *Cckar* expression was detected between groups at P21, further confirming that CCK receptor expression does not drive the differential satiation responses between groups. Genes related to inflammation were well represented among the DEGs, including higher expression of *Angptl2*, *Angptl4*, *Gbp2*, *Gbp6*, *Ifit3*, and *Il1rap*. Diets high in fats have been demonstrated to induce inflammation in the brain of adult humans and rodents, as evidenced by increased microglial infiltration in the hypothalamus and other brain regions^36^. Additionally, several genes involved in modulating neuronal excitability were differentially expressed, including increased *Atp1a2* (an ion pump that modulates neuronal excitability), decreased *Fxyd7* (which limits neuronal excitability), and decreased *Camk2n2* (which limits Camk2 dependent activity) in HFD^EARLY^ mice (Figure 3D).

Using the gene sets described in Supplemental Figure 3, we assessed whether expression of genes in key domains related to developmental processes differed between control and HFD^EARLY^ mice at P21. Many of the domains exhibited non-significant differences in summed expression including cell cycle/proliferation, myelin, and postsynaptic density and ligand gated ion channels, among others (data not shown). However, two domains did exhibit significantly reduced summed expression in HFD^EARLY^ mice: cytoskeletal organization, and synaptic vesicles. Consistent with the results from bulk sequencing, immunostaining for synaptophysin (a membrane glycoprotein enriched in synaptic vesicles and widely used as a marker of presynaptic terminals) demonstrates that at P21 HFD^EARLY^ mice exhibited a significantly lower density of synaptophysin-labelled puncta in the nucleus of the solitary tract (NTS; Figure 3F), where vagal sensory neurons project.

### Early Life HFD does not alter adult CCK-dependent satiation responses

To determine whether early life diet alters CCK-dependent satiation signaling in adults, mice were also tested at P120. Following saline injection, there was no difference in consumption between control and HFD^EARLY^ mice. Additionally, following CCK injection, there was no difference in consumption between control and HFD^EARLY^ mice (Figure 4K-N). The data demonstrate that CCK-dependent satiation signaling remains intact in adult HFD^EARLY^ mice. Despite normal satiation responses, a modest increase in body weight was observed in HFD^EARLY^ mice of both sexes at P120, however this was accompanied by no significant differences in adiposity between sex-matched groups (Figure 4A-J). A small, but statistically significant difference in lean mass was observed in HFD^EARLY^ males. Home cage chow consumption was measured, revealing no significant difference in average daily consumption (Figure 4O).

**Figure 4.**
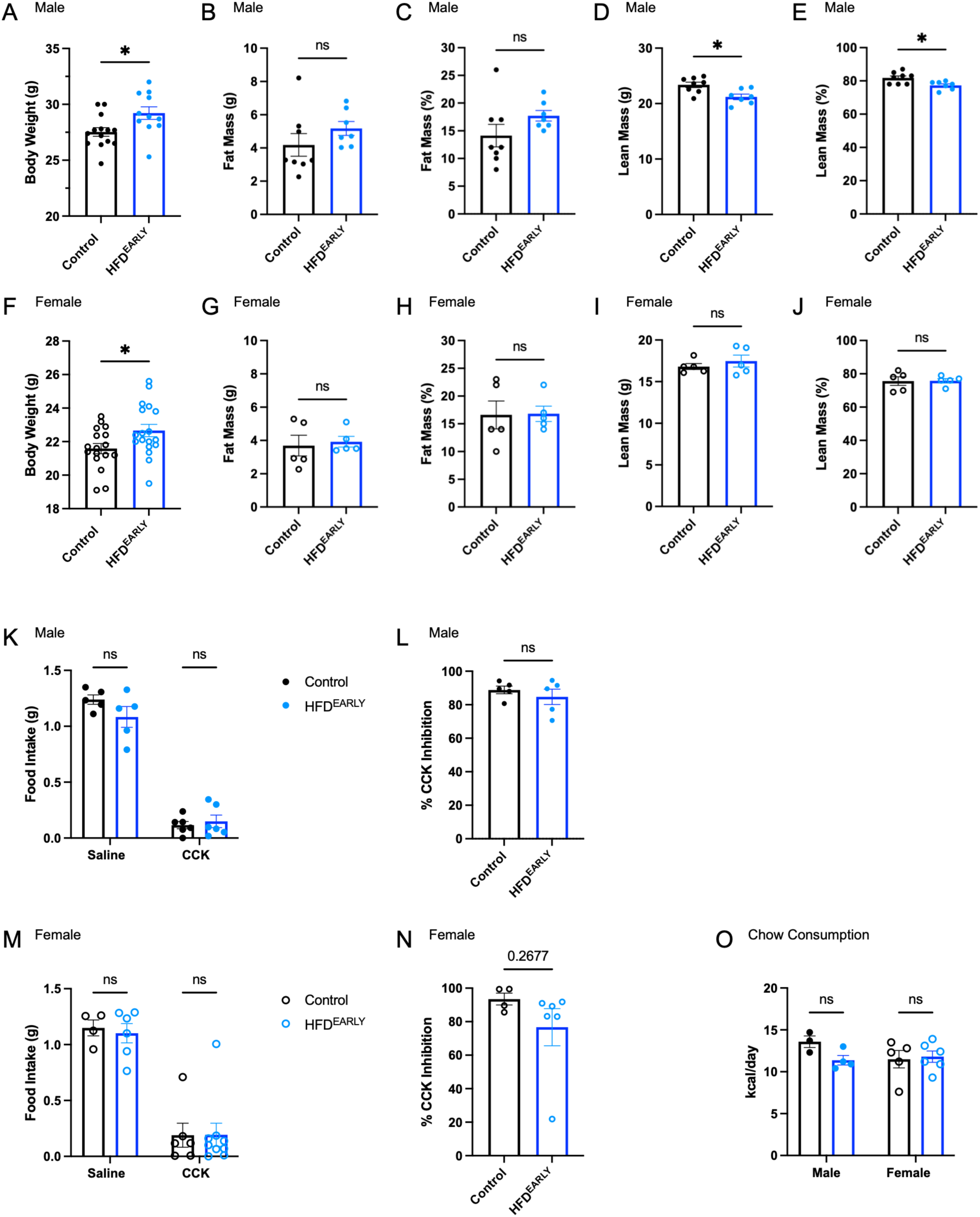
Early life HFD does not alter adult CCK-dependent satiation responses but produces modest changes in adult body composition. A-E) Body weight, fat mass (g), fat mass (%), lean mass (g), and lean mass (%) in male control and HFD^EARLY^ mice at P120. F-J) Body weight, fat mass (g), fat mass (%), lean mass (g), and lean mass (%) in female control and HFD^EARLY^ mice at P120. K) Food (HFD) intake in adult males at P120 following saline or CCK injection. L) Percent feeding inhibition in adult males following HFD testing at P120. M) Food (HFD) intake in adult females at P120 following saline or CCK injection. N) Percent feeding inhibition in adult females following HFD testing at P120. O) Average daily home-cage chow consumption in kcal/day in male and female control and HFD^EARLY^ mice at P120. A two-tailed unpaired t-test was used for body composition and percent feeding inhibition comparisons. A two-way mixed-effects ANOVA was used to analyze differences in response to saline or CCK between groups. * p<0.05; ns, not significant.

### Early life HFD exposure increases susceptibility to overconsumption of a calorie-dense liquid diet in adulthood

The observation that standard chow intake did not differ between control and HFD^EARLY^ mice at P120 (Figure 4O) raises the question of whether the modest increase in body weight observed in these animals reflects latent alterations in intake that are intensified under conditions of heightened dietary challenge. To address this, adult control and HFD^EARLY^ mice (P120) were provided *ad libitum* access to a calorie-dense liquid nutritional supplement (BOOST® Very High Calorie, 2.24 kcal/mL) alongside standard chow and water for 7 days. Daily chow intake did not differ between groups in either sex across the challenge period (Figure 5B and I). However, female HFD^EARLY^ mice consumed significantly more of the calorie-dense liquid diet and more calories overall than control females (Figure 5I and J). Female HFD^EARLY^ mice also exhibited a trend toward greater body weight gain over the 7-day period (Figure 5K; p=0.075), and a significant increase in fat mass and decrease in lean mass relative to controls at the end of the challenge (Figure 5L and M). A similar trend was observed in the males that did not reach statistical significance. Together, these findings suggest that early life HFD exposure produces a lasting increase in susceptibility to overconsumption of calorie-dense foods, an effect that becomes apparent under dietary conditions that model modern energy-dense environments associated with obesity risk.

**Figure 5.**
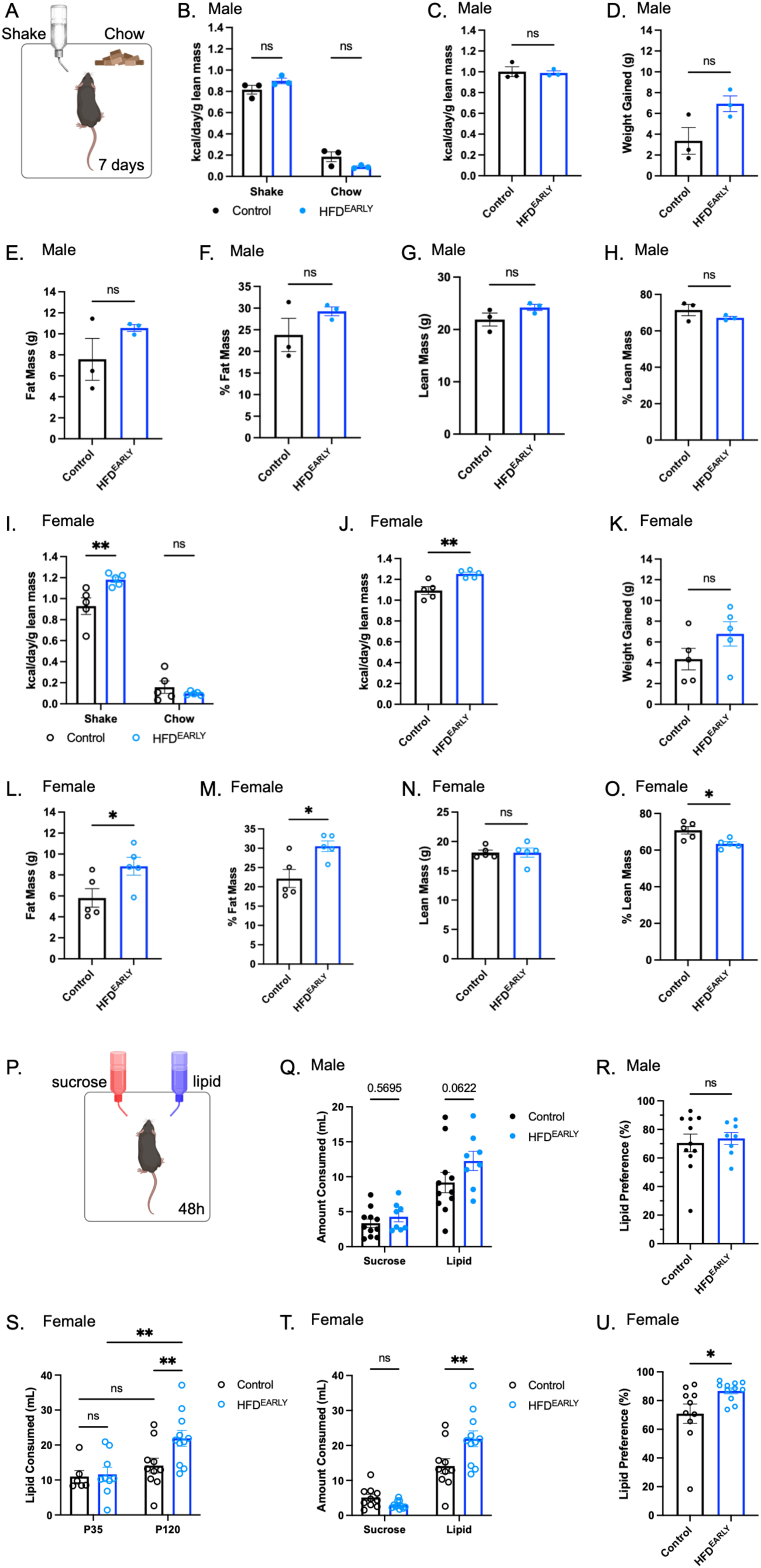
Female mice overconsume a calorie-dense liquid diet in adulthood and overconsume and prefer lipid in a two-bottle preference test. A) Schematic of the calorie-dense liquid diet challenge. B-D) Daily intake of BOOST Very High Calorie shake and chow (kcal/day/g lean mass) and body weight gained (g) in male control and HFD^EARLY^ mice over the 7-day challenge (∼P120) (n=3). E-H) Fat mass (g), fat mass (%), lean mass (g), and lean mass (%) in males at the conclusion of the challenge. I-K) Daily intake of BOOST shake and chow (kcal/day/g lean mass) and body weight gained (g) in female control and HFD^EARLY^ mice over the 7-day challenge (n=5). L-O) Fat mass (g), fat mass (%), lean mass (g), and lean mass (%) in females at the conclusion of the challenge. P) Schematic of the two-bottle preference test (48 hours). Q) Amount of sucrose and intralipid consumed by male control and HFD^EARLY^ mice at P120 during the 48-hour test (n=8-11). R) Lipid preference ratio in males. S) Comparison of the amount of lipid consumed by each group at P35 and P120 in females, demonstrating that the overconsumption phenotype observed in HFD^EARLY^ females does not emerge until adulthood. T) Amount of sucrose and intralipid consumed by female control and HFD^EARLY^ mice at P120 during the 48-hour test (n=10-11). U) Lipid preference ratio in females. A two-tailed unpaired t-test was used to analyze body composition, body weight gained, and lipid preference. A two-way mixed-effects ANOVA was used to analyze daily intake across the 7-day challenge and differences in amount consumed in the two-bottle preference test (* p<0.05, ** p<0.01). ns, not significant.

### Adult HFD^EARLY^ mice overconsume lipids

Ingestion of fats and sugars engages vagal sensory neurons that connect to reward responsive pathways, reinforcing consumption^1–3,6^. To determine whether early life HFD alters consumption of fats and sugars known to engage these pathways, a two-bottle preference test was used. On P35, control and HFD^EARLY^ mice were provided a choice between a solution of 0.6% lipid (fat emulsion; intralipid) or 1.5% sucrose (Figure 6A). Solutions in each bottle were matched for caloric content at low calorie-density (0.06kcal/ml), designed to test nutrient preference while minimizing the influence of calorie-driven effects. Based on the two-bottle preference test, all mice preferred lipid over sucrose, suggesting that the circuits required to form a nutrient-specific preference are functional at this age (Supplemental Figure 4). No significant differences were observed in the amount of sucrose or lipid consumed between control and HFD^EARLY^ mice of either sex at P35 (Supplemental Figure 4). By contrast, when the 2-bottle preference test was applied at a later age, P120, HFD^EARLY^ females consumed significantly more (∼50%) intralipid than sex-matched controls (Figure 5T). HFD^EARLY^ females also exhibited a greater preference for lipid than control females (Figure 5U). HFD^EARLY^ males also exhibited a trend in increased consumption of lipid that did not reach statistical significance (Figure 5Q and R). Together, the data suggest that early life HFD exposure induces overconsumption of fats in adulthood, with sex modulating the effect size.

**Figure 6.**
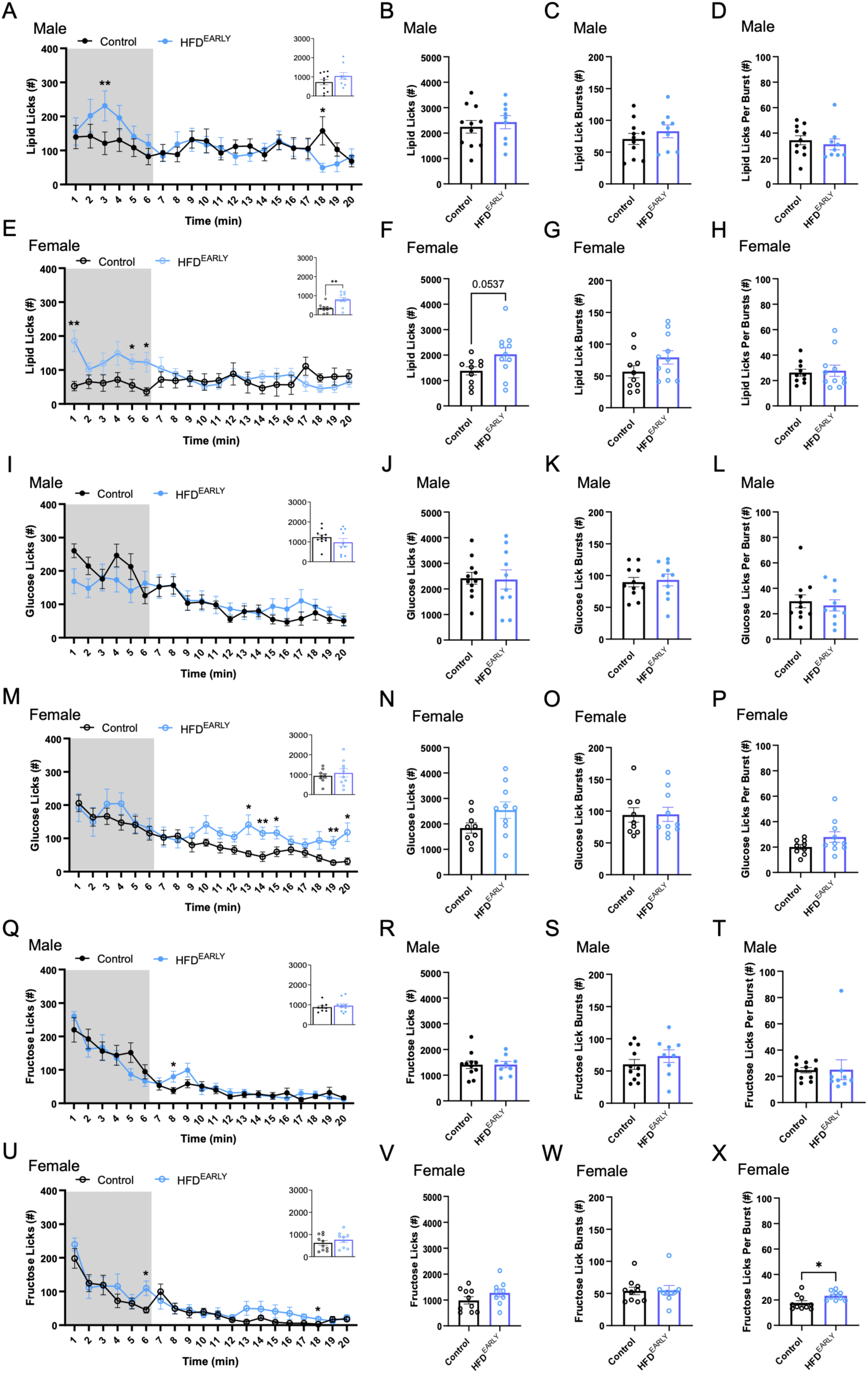
Exposure to high fat diet early in life changes nutrient consumption patterns in females. A-D) Licking rate per minute, total licks, number of licking bursts, and average licking burst size across the single-bottle intake session with 5% Intralipid for HFD^EARLY^ and control males (n=9-11). Inset in A shows the total number of licks within the first 6 minutes of the test (appetition phase, shaded). E-H) Licking rate per minute, total licks, number of licking bursts, and average licking burst size across the single-bottle intake session with 5% Intralipid for HFD^EARLY^ and control females (n=10-11). Inset in E shows total licks during the appetition phase. I-L) Licking rate per minute, total licks, number of licking bursts, and average licking burst size across the single-bottle intake session with 10% glucose for HFD^EARLY^ and control males (n=10-11). Inset in I shows total licks during the appetition phase. M-P) Licking rate per minute, total licks, number of licking bursts, and average licking burst size across the single-bottle intake session with 10% glucose for HFD^EARLY^ and control females (n=9-10). Inset in M shows total licks during the appetition phase. Q-T) Licking rate per minute, total licks, number of licking bursts, and average licking burst size across the single-bottle intake session with 10% fructose for HFD^EARLY^ and control males (n=9-10). Inset in Q shows total licks during the appetition phase. U-X) Licking rate per minute, total licks, number of licking bursts, and average licking burst size across the single-bottle intake session with 10% fructose for HFD^EARLY^ and control females (n=9-10). Inset in U shows total licks during the appetition phase. Line graphs show mean +/-SEM for each group; bar graphs show mean +/-SEM with individual data points. The shaded portion of panels A, E, I, M, Q, and U highlights the early-meal appetition phase. A repeated-measures two-way ANOVA with post hoc Tukey correction was used to analyze differences in licks per minute between groups (* p<0.05, ** p<0.01). A two-tailed unpaired t-test was used to analyze cumulative licks, burst number, and licks per burst between groups (* p<0.05, ** p<0.01).

### Early life HFD exposure drives nutrient-specific changes in early appetition and satiation

To further investigate differences in consumption in response to nutrient-specific stimuli, we performed a series of single-nutrient acceptance tests and measured the microstructural patterns of intake (Figure 6A-H). In line with the results of the two-bottle preference test, male control and HFD^EARLY^ mice consumed similar overall amounts of lipid over the 20-minute test (Figure 6A-B). The inset in figure 6A represents the total number of licks during the first 6 minutes of the test, when the signals that rapidly reinforce ingestion are engaged (Figure 6A inset)^37,38^. HFD^EARLY^ male mice did have a transient spike in licking for lipid early in the session (minute 3), however no significant difference was identified in the total lipid licks within the first 6 minutes (Figure 6A inset). No other differences in licking patterns were observed in males (Figure 6C-D). Responses to lipid differed by sex where females from the HFD^EARLY^ group consumed more of the 5% lipid solution than control females (Figure 6E-F), and this was driven by significantly elevated intake during the early phase of the test (Figure 6E inset). The overall licking patterns were otherwise similar among the two groups (Figure 6G-H).

Intake is primarily controlled by oral, proximal gut sensory signals, or their interaction during the initial part of a meal^37,38^. Therefore, a series of brief access taste tests was used to parse the contribution of oral sensing to nutritive reward (Figure 7). In these tests, mice sample a range of stimulus concentrations in 10-sec trials, minimizing postingestive effects on intake in each trial, and focusing on orosensory-driven behavior. HFD^EARLY^ and control mice licked in similar concentration-dependent manners for lipid, except at the highest concentrations of lipid tested (Figure 7B and D). Both HFD^EARLY^ male and female mice licked more avidly than sex-matched controls for the highest concentration of lipid tested. HFD^EARLY^ and control mice initiated a similar number of trials to obtain lipid solutions (Figure 7C and E), suggesting that the history with HFD early in life did not impact orosensory-mediated appetitive responding for fat. The collective results suggest that the attraction to the ‘taste’ of fat is amplified in both sexes at high concentrations as a result of the early life HFD.

**Figure 7.**
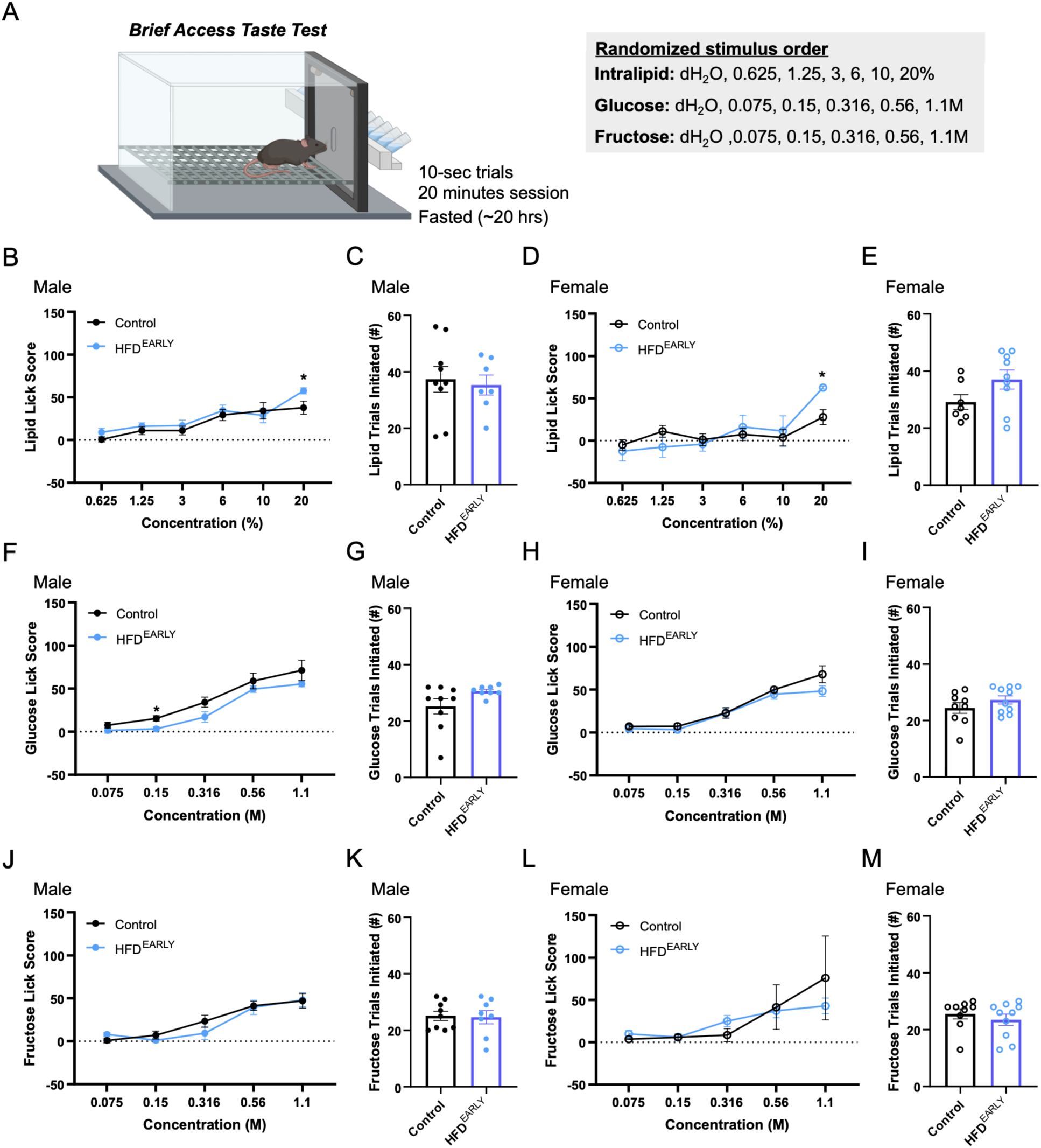
Early life HFD exposure increases licking avidity for high concentrations of lipid. A) Schematic of the brief access taste test chamber and protocol. Stimulus concentrations tested: Intralipid 0.625%, 1.25%, 3%, 6%, 10%, 20%; Glucose 0.075, 0.15, 0.316, 0.56, 1.1 M; Fructose 0.075, 0.15, 0.316, 0.56, 1.1 M. Lick score = mean licks per concentration minus mean water licks per session. B-C) Lipid lick score and total lipid trials initiated for HFD^EARLY^ and control males (n=7-9). D-E) Lipid lick score and total lipid trials initiated for HFD^EARLY^ and control females (n=7-9). F-G) Glucose lick score and total glucose trials initiated for HFD^EARLY^ and control males (n=8-9). H-I) Glucose lick score and total glucose trials initiated for HFD^EARLY^ and control females (n=9-10). J-K) Fructose lick score and total fructose trials initiated for HFD^EARLY^ and control males (n=9-10). L-M) Fructose lick score and total fructose trials initiated for HFD^EARLY^ and control females (n=8-9). Line graphs show mean +/-SEM. A repeated-measures two-way ANOVA with post hoc Tukey correction was used to analyze differences in lick score across concentrations between groups (* p<0.05).

To similarly understand whether responses to ingested sugars were impacted, brief access taste tests (Figure 7F-M) and single nutrient acceptance tests (Figure 6I-X) were conducted, with glucose and fructose presented in separate sessions. Although both solutions taste sweet, glucose has greater reinforcing efficacy than fructose in C57BL6/J mice^39^. In general, intake of each sugar was comparable across early life diet groups, for both the males and the females (Figure 6I-X). However, HFD^EARLY^ females maintained a consistent level of glucose intake across the entire 20-minute session, whereas the control females showed a normal cessation of intake as the session progressed (Figure 6M). Taste-motivated responses to each sugar were largely unaffected by early life diet, with the exception of a blunted sensitivity to perithreshold glucose in male HFD^EARLY^ mice (Figure 7F). Therefore, the persistent ingestion of 10% glucose in the single bottle acceptance tests likely reflects an impaired satiation mechanism specific to glucose or a delayed reward signal. Notably, while total licks for fructose did not differ between groups, HFD^EARLY^ females performed more licks per burst, which may suggest an increased sensitivity to intake-promoting properties of fructose. Taken together, these findings indicate that early life exposure to HFD alters distinct aspects of satiation and appetitive reward for sugars and fats.

## Discussion

The Developmental Origins of Health and Disease (DOHaD) framework posits that the nutritional environment experienced during critical periods of early development exerts lasting effects on physiology and disease risk that persist into adulthood. A substantial body of work in rodent models has established that perinatal overnutrition — achieved through high-fat diet feeding during gestation and lactation, or through litter size manipulation — programs lasting increases in adiposity, insulin resistance, and hyperphagia in offspring^40–47^. These effects arise, in part, through persistent changes in the development of hypothalamic circuits regulating energy balance, including alterations in leptin signaling, orexigenic neuropeptide expression, and the formation of melanocortin projections as well as alterations in mesolimbic dopamine signaling^25,48–52^. The present findings extend the DOHaD framework to include vagal interoceptive circuits as a target for early nutritional programming. We demonstrate that CCK-dependent satiation matures precociously following early-life HFD exposure through putative changes in gene expression altering membrane polarization. Critically, this early maturational and transcriptional shift is accompanied by lasting consequences: adult HFD^EARLY^ mice show sex-specific alterations in appetitive behavior, overconsumption of palatable, energy-dense foods, and resulting increases in adiposity. Together, these findings suggest that the early nutritional environment shapes vagal circuit development in ways that are mechanistically distinct from the effects of HFD exposure in the mature nervous system.

### The effects of HFD on CCK-dependent satiation signaling depend on the timing of the dietary exposure

Several studies completed in adult rodents and humans demonstrate that extended periods of exposure (8+ weeks) to high-fat or calorie dense foods can impair vagally-mediated satiation^32,33,53^. We replicate that effect here, demonstrating that modest impairments in CCK-dependent feeding inhibition can be observed after only 3 weeks on a HFD in adult mice. Furthermore, we demonstrate that HFD exposure in early life prior to weaning has the opposite effect on developing vagal circuits, promoting enhanced, more mature patterns of CCK-dependent satiation behaviors by P35. Together, these data establish early life as a critical period for vagal sensory circuit development, and demonstrate that the mechanisms driving alterations in vagal function differ fundamentally depending on the timing of dietary exposure.

The mechanistic basis for this timing-dependent divergence in phenotype is likely rooted in the distinct biological states of the vagal circuit at the time of dietary exposure. Published work in rats suggests that the mechanism underlying decreased CCK sensitivity in adult HFD-fed animals involves obesity-induced leptin resistance and a resulting inhibition of CCK signaling in VSNs^32,53^. HFD^EARLY^ mice did exhibit greater adiposity at P35, but did not weigh significantly more at the time of testing, making it unlikely that they had achieved a degree of leptin resistance comparable to that observed in the rats described in the aforementioned study. Other studies have examined the effects of perinatal HFD in the absence of obesity on the dorsal motor nucleus of the vagus, demonstrating that perinatal HFD reduces the excitability of descending motor neurons as well as the ability of their afferents to respond to CCK, implicating effects on neural excitability in connected circuits^54^. Consistent with this, our bulk RNA sequencing data suggest that differential expression of genes associated with increased VSN excitability (*Atp1a2, Fxyd7,* and *Camk2n2*) likely drive early maturation of these neurons and their associated behaviors. The accompanying reduction in synaptophysin-labeled puncta density in the NTS may reflect activity-dependent synaptic pruning— a process by which heightened afferent activity drives elimination of excess synaptic contacts as circuits refine and stabilize. Such pruning is well documented in other developing sensory systems and is generally associated with, rather than opposed to, functional maturation. It is important to note, however, that synaptophysin labels synaptic vesicle proteins broadly and does not distinguish between excitatory and inhibitory terminals, nor between vagal afferent inputs and local interneuron synapses within the NTS. Whether the reduction in puncta density reflects remodeling of vagal afferent terminals specifically, or changes in the broader synaptic milieu of the NTS, remains to be determined.

A parsimonious upstream explanation for the precocious maturation of CCK-dependent satiation in HFD^EARLY^ mice is that elevated CCK secretion during the early-life dietary manipulation itself drives accelerated development of the vagal circuits that mediate its effects. CCK is released from intestinal enteroendocrine cells in proportion to luminal fat content^55^, and the elevated lipid content of milk from HFD-fed dams would be expected to produce sustained increases in CCK secretion in nursing pups^34,35,56,57^. Additionally, as pups begin consuming solid food during the complementary feeding period, direct exposure to HFD pellets would provide a second route of heightened CCK stimulation. Repeated or sustained CCK receptor activation during a period of active synaptogenesis and circuit maturation could plausibly accelerate the functional maturation of satiation responses independently of changes in *Cckar* transcript levels, which did not differ between dietary groups at any age examined. Instead, the transcriptomic evidence of altered VSN excitability observed here may reflect downstream consequences of chronic CCK receptor activation on the intrinsic properties of vagal sensory neurons. While we did not directly measure CCK secretion or VSN functional activity in these animals, this hypothesis is consistent with the well-established relationship between dietary fat intake and CCK release. Direct measurement of CCK secretion in HFD^EARLY^ pups, dissociation of the contributions of the milk-borne vs. direct dietary exposure periods, and direct electrophysiological or calcium imaging measures of VSN excitability will be important goals for future work.

### Effects of early life HFD on taste driven motivation, particularly for concentrated lipid

The peripheral end organ and first order central relay of the gustatory system mature during the late prenatal and early postnatal period, with continued refinement through adolescence^58–62^. The development of the gustatory system is shaped by diet, both via maternal influence and through direct consumptive experience with foods early in life^63–65^. In light of this, we used brief access taste tests to determine whether the changes in ingestive behaviors for lipids and sugars were directly related to altered taste function in HFD^EARLY^ mice. We found that HFD^EARLY^ males and females licked comparably to their control counterparts at lower lipid concentrations but licked more avidly for 20% lipid. HFD^EARLY^ males also had a reduced sensitivity to glucose at a low, perithreshold concentration. Prior studies have shown mixed results on the effects of maternal HFD (including during gestation and/or lactation) on nutrient taste in rats and mice^66–69^. This variability may reflect differences in the timing and degree of dietary exposure, or in the age and method used to assess taste sensitivity^66^. Gestation and lactation HFD have also been shown to alter taste receptor profiles in the offspring^68,69^. Whether the heightened responsiveness to concentrated lipid and blunted sensitivity to glucose observed here are driven by programmatic shifts in taste and/or somatosensory receptors in the oral cavity or in the higher-order brain systems that process orosensory information remains to be determined. Notably, the amplified orosensory response to concentrated lipid in both sexes likely contributes to the exaggerated early-meal appetition phenotype observed in female HFD^EARLY^ mice during single-bottle acceptance testing, suggesting that orosensory and post-ingestive mechanisms may act in concert to drive the adult overconsumption phenotype.

### Durable consequences of early life HFD consumption on post-ingestive reward behaviors

The two-bottle preference and lickometer data reveal that early-life HFD produces lasting, nutrient- and sex-specific alterations in adult consumptive behavior. HFD^EARLY^ females overconsume and prefer lipid over sucrose in the two-bottle test. Because fat sensation is conveyed to the brain specifically via vagal afferents — unlike sugar, which engages both vagal and spinal pathways^70^ — the selective lipid overconsumption points to altered signaling via vagal afferent circuits. Microstructural analysis further revealed that HFD^EARLY^ females showed an exaggerated appetition response when consuming lipid, and a protracted intake pattern when consuming glucose, consistent with impaired glucose-driven satiation or delayed postingestive reinforcement. As fructose, which has reduced postingestive rewarding properties compared to glucose and lipid in C57BL/6J mice^30,31^, did not produce a similar late-meal response, we favor the latter interpretation. More studies will be needed to explicitly test this. These effects were not overt in males, whose overconsumption of lipid did not reach significance and who showed no clear shift in nutrient preference.

The female specificity of these behavioral phenotypes likely reflects sex differences in the recruitment of downstream reward and motivational circuits. Few genes are differentially expressed in VSNs from males compared to females, pointing to sexually dimorphic circuitry downstream of the vagus rather than at the level of primary afferents as the source of the behavioral divergence. Recently published work using the same maternal HFD model supports this interpretation, reporting changes in dopaminergic circuitry alongside increased sucrose intake and weight gain specifically in female mice ^25,51^. Whether these dopaminergic impairments are mechanistically driven by enhanced excitatory drive from VSNs, and why males appear protected from the lasting behavioral consequences of early-life HFD, remain important open questions.

Considered together, the appetitive phenotypes offer a potential explanation for a seemingly paradoxical finding: despite normal home-cage chow intake at P120, HFD^EARLY^ mice show a modest but persistent increase in body weight. The calorie-dense liquid diet challenge directly addressed this apparent contradiction. Female HFD^EARLY^ mice selectively overconsume the liquid supplement – but not chow – resulting in increased fat mass, with a trend toward greater body weight gain (p=0.075), with only 7 days of access. This pattern is consistent with a context-dependent susceptibility to overconsumption of palatable, energy-dense foods that is dissociable from baseline appetite chow consumption. Gonadal hormones such as estrogen, which modulate both energy balance and sensitivity to palatable food reward, may further amplify this female-specific vulnerability^71–73^. Together, these findings suggest that early-life dietary programming produces a latent susceptibility to overconsumption that becomes apparent under conditions that more closely resemble the energy-dense food environments associated with human obesity risk.

### Limitations

Several limitations of the present study warrant consideration. First, the early life dietary manipulation employed a purified high-fat diet (D12451) that differs from standard colony chow not only in fat content but also in carbohydrate source, protein source, and fiber composition. Although D12451 is among the most widely used rodent HFD formulations in the field, these compositional differences make it difficult to attribute the observed effects on vagal circuit maturation exclusively to elevated dietary fat. Future studies employing isocaloric diets that vary a single macronutrient will be necessary to isolate the specific dietary component responsible for programming vagal satiation circuits during early life. Second, the present model does not permit dissociation of the relative contributions of maternal milk composition and direct pup consumption of HFD during the complementary feeding period to the observed phenotypes. Dam HFD consumption alters milk lipid content, exposing pups to elevated dietary fat from birth; however, pups also begin directly consuming solid HFD during the second and third weeks of life. Whether the critical window for programming vagal satiation circuits falls within the nursing period, the complementary feeding period, or both remains an important open question, and will be addressed in future work. Third, HFD consumption is known to alter gut microbiome composition, and the microbiome exerts bidirectional influence on vagal afferent signaling through the production of short-chain fatty acids, bile acids, and other gut-derived metabolites. We cannot exclude the possibility that microbiome differences between dietary groups contribute to the altered VSN gene expression and satiation phenotypes observed here. Future studies employing germ-free or antibiotic-treated animals will be necessary to isolate the contribution of microbiome-dependent signaling to the effects of early life HFD on vagal circuit development.

## Conclusion

In conclusion, our findings establish early life as a critical period for the development and functional tuning of vagal satiation circuits, with lasting consequences for adult feeding behavior and metabolic health. Using a mouse model of preweaning HFD exposure, we demonstrate that early dietary experience can accelerate the maturation of CCK-dependent satiation responses, an effect that is mechanistically distinct from the impairments produced by adult-onset HFD exposure. This developmental programming is accompanied by altered expression of genes controlling the intrinsic excitability of VSNs, suggesting that transcriptional remodeling of vagal sensory neurons contributes to the observed phenotype. Furthermore, early-life HFD produces lasting, sex-specific alterations in appetitive behavior and susceptibility to overconsumption of palatable energy-dense foods, pointing to the recruitment of sexually dimorphic downstream circuits, particularly dopaminergic pathways, as a key determinant of adult metabolic outcomes. These results extend the DOHaD framework to vagal interoceptive circuits and contribute to a growing body of evidence that early nutritional environments have enduring, circuit-specific impacts on the regulation of feeding behavior. Future work will be essential to identify the cellular and molecular mechanisms driving these nutrient-and sex-specific outcomes, measure VSN functional activity directly, and explore whether interventions targeting the early nutritional environment could mitigate their lasting effects on metabolic health.

## Acknowledgements

The image acquisition work was performed in the Cellular Imaging Core at The Saban Research Institute (TSRI), Children’s Hospital Los Angeles (CHLA). The RNA Sequencing work was performed in the Spatial Biology and Genomics Core, CHLA. The body composition measurements were performed in the Metabolic Phenotyping Core at CHLA.

We thank Samuel Egboh, Rachel Moreno, and Kimberly Ruvalcaba from the Samuels Family Los Angeles High School Internship Program (LA-HIP) and Chuck Lorre Research Scholars Program for their work quantifying in situ hybridization data.

## Author Contributions

MM Performed Research, Analyzed Data, Wrote the Paper, Revised the Paper; CR Performed Research, Analyzed Data, Wrote the Paper, Revised the Paper; JP Analyzed the Data; LS Designed Research, Wrote the Paper, Revised the Paper; AK Designed and Performed Research, Analyzed Data, Wrote the Paper, Revised the Paper

## STAR METHODS

## EXPERIMENTAL MODEL AND STUDY PARTICIPANT DETAILS

### Animals

Animal husbandry and experimental procedures were conducted in accordance with the guidelines of the Institutional Animal Care and Use Committee of The Saban Research Institute of Children’s Hospital Los Angeles (CHLA) and the University of Southern California (USC). C57BL/6J mice [strain 000664; RRID: IMSR_JAX:000664] were obtained from The Jackson Laboratory and bred in the animal care facilities at CHLA and USC. All mice were wild-type and were treatment-naive unless otherwise stated. Mice were housed in the vivarium on a 13 h light/11 h dark cycle at CHLA or a 12 h light/12 h dark cycle at USC, with housing maintained at 22°C.

### Early-life high-fat diet paradigm

Pregnant dams were maintained on standard chow (13% kcal from fat, 62% kcal from carbohydrate [3% from sucrose], 24% kcal from protein; PicoLab Rodent Diet 20, catalog #5053, LabDiet) until parturition. Because litter size influences early ingestive experience and weight gain^74,75^, litters were culled to 6-8 pups on postnatal day (P) 0 with care taken to maintain a balanced sex ratio. Litters meeting the size criteria were randomly assigned to either remain on standard chow (Control) or receive a high-fat diet (HFD^EARLY^; 45% kcal from fat, 35% kcal from carbohydrate [17% from sucrose], 20% kcal from protein; Irradiated Rodent Diet with 45 kcal% Fat, catalog #D12451i, Research Diets, Inc.) from P0 until weaning at P21. During this period, food pellets were also placed on the cage floor to allow pups access during the complementary feeding period as they began to consume solid foods. All offspring were maintained on standard chow from P21 until the end of the study unless otherwise specified.

### Adult high-fat diet paradigm

C57BL/6J mice were raised on standard chow. At P85, mice were randomized into groups maintained on either standard chow (Control; PicoLab Rodent Diet 20; catalog #5053, LabDiet) or a high-fat diet (HFD^LATE^; catalog #D12451i, Research Diets, Inc.) for three weeks. This duration was selected to match the three-week period of early-life HFD exposure in the HFD^EARLY^ paradigm, thereby allowing direct comparison of the effects of HFD exposure across developmental stages. After the three-week experimental period, all mice were maintained on standard chow until the end of the study.

### Experimental design and blinding

No additional randomization or stratification procedures were used beyond random assignment of qualifying litters to dietary groups and randomization of adult mice to dietary groups. Behavioral experiments were conducted and analyzed with investigators blinded to experimental group. Slides were coded so that investigators were blinded to experimental group during imaging and image analysis. Individual inclusion and exclusion criteria are described in the corresponding experimental procedures below.

## METHOD DETAILS

### Body weight and body composition measures

Body weights were recorded weekly during the dietary manipulation period for both the early-life and adult diet paradigms. Body composition was measured at P35 and P120 using an EchoMRI-100H Body Composition Analyzer. Additional body composition measurements were taken at the conclusion of the calorie-dense liquid diet challenge.

### Assay of CCK-dependent satiation

Mice were single-housed and fasted overnight for approximately 16 h before each test day, with food removed at approximately 5:00 PM and testing conducted at approximately 9:00 AM during the light phase. On test days, mice received an intraperitoneal (i.p.) injection of either sterile saline or cholecystokinin-8 (CCK-8; Cholecystokinin-octapeptide, sulfated; catalog #H-2080.0001BA; Avantor, Inc.) at a dose of 10 µg/kg body weight in a volume of 1 µL/g body weight. Immediately following injection, mice were presented with a pre-weighed pellet of either standard chow or high-fat diet. Food consumption was recorded at 30 min and 1 h post-injection.

All mice received a habituation saline injection at least 3 days before the first experimental trial to acclimate them to the injection procedure; data from this habituation session were not used in analyses. Subsequently, each mouse served as its own control by receiving both a saline injection and a CCK injection on separate test days spaced at least 3 days apart. CCK injections were timed to the target age of the assay +/- 2 days. Saline injections always followed 3 days after the CCK-injection test. Percent feeding inhibition was calculated for each animal by comparing food intake following CCK injection with intake following the post-habituation saline injection. Mice were tested at P35, P45, P60, and P120 for the developmental series and at P35 and P120 for dietary group comparisons. Given the protracted fasting period that is standard for this test, earlier ages were not evaluated due to health concerns.

### Calorie-dense liquid diet challenge

Male and female control and HFD^EARLY^ mice at approximately P120 (±2 weeks) were individually housed and provided *ad libitum* access to a bottle containing BOOST Very High Calorie nutritional drink (Very Vanilla flavor; 2.24 kcal/mL; Nestlé) in addition to standard chow and water for 7 days. Bottles containing liquid diet and chow were weighed every 24 h to assess daily intake. Body weight was measured on days 0, 3, and 7 of the challenge. Body composition was measured at the beginning and conclusion of the challenge (days 0 and 7, respectively).

### Two-bottle preference tests

Two-bottle preference testing was conducted at P20-25, P30-35, P40-45, P50-55, and approximately P120 (±2 weeks) to determine the age at which post-ingestive appetitive responding matured. The age ranges reflect the fact that the test takes 5 days to run and occurred within each age range ±2 days. Mice were housed with a sex-matched littermate (2 per cage) and given 24 h to acclimate to two drinking bottles containing water. On the first training day, mice received one bottle containing water and one containing 10% glucose in water. On the second training day, mice received one bottle containing water and one containing 10% fructose in water. On the test day, mice received one bottle containing 10% glucose and one containing 10% fructose, which were calorie matched (0.34 kcal/mL). Bottle location was reversed from the training days. Fluid intake was measured by weighing each bottle at the start of the test and every 24 h thereafter. Mice had *ad libitum* access to standard chow throughout the test.

A separate two-bottle test was conducted in control and HFD^EARLY^ offspring at P35 and P120 to examine sensitivity to fats compared with sugars. Mice were individually housed and given 24 h to acclimate to two drinking bottles containing water. At the start of the test, mice were provided one bottle containing 0.6% lipid (Intralipid, catalog #831818311, Fresenius Kabi USA) in water and one bottle containing 1.5% sucrose (catalog #84097, Sigma-Aldrich) in water. Solutions were calorie matched at 0.06 kcal/mL. The test period lasted 48 h, and bottle positions were switched after the first 24 h to control for side preference. Fluid intake was measured by weighing each bottle at the start of the test and every 24 h thereafter. Mice had *ad libitum* access to standard chow throughout the test.

### Nutrient acceptance and licking microstructure

Consumption patterns of fat (5% lipid; Intralipid) and calorically matched representative sugars (10% glucose and 10% fructose) were measured in a lickometer (Davis Rig; Med Associates Inc., Fairfax, VT) in a subset of mice at P120-160 from each treatment group. Mice were initially water deprived for approximately 20 h and trained to lick for water across two 20-min sessions conducted on consecutive days. Successful training was defined as completing at least 1,000 licks on the second training day. Water was returned to the home cage 30 min after the criterion was met. Only animals meeting this criterion were tested.

Following training, mice were given one rest day, and chow was removed for approximately 20 h before each test session. Each test session was separated by at least 2 days. For testing, mice were given access to a single nutrient in the lickometer for 20 min, and time-stamped licks were recorded for microstructural analyses. Chow was returned 30 min after each test. Only one nutrient was tested per day. All mice were first tested with lipid, and the order of glucose and fructose testing was counterbalanced across mice. Nutrients were prepared in deionized water on the day of testing.

Total licks, burst size (number of licks per burst), and total number of bursts were analyzed using LickCalc software (https://lickcalc.uit.no/). Licking bursts were defined as a run of at least 3 consecutive licks separated by less than a 1-s pause.

### Brief access taste tests

Brief access taste tests were conducted in the Davis Rig following completion of all nutrient acceptance tests to measure licking responses motivated by the taste of nutrients apart from post-ingestive effects. Following a 20-h food deprivation period, mice were presented with various concentrations of one tastant and water in successive 10-s trials for 20 min. Mice were permitted to initiate an unlimited number of trials during each session. Lipid concentrations were 0.625%, 1.25%, 3%, 6%, and 10% in the Methods protocol. Glucose and fructose concentrations were 0.075, 0.15, 0.316, 0.56, and 1 M. Trials were organized into blocks in which tastant concentrations were randomly ordered, with no concentration repeated within a block. Each trial lasted 10 s from the first lick.

Lick score was calculated as the mean licks for each tastant concentration minus the mean licks for water. Only mice that completed at least two trials per solution were included in the analysis, and the number of licks for each solution was averaged across trials.

### RNA sequencing

Nodose ganglia were dissected from male P21 and P35 mice with the aid of a dissecting microscope. Left and right nodose ganglia were combined for each animal, flash-frozen over powdered dry ice, and stored at -80°C until further processing. RNA was isolated using the RNeasy Micro Kit (QIAGEN, catalog #74004) according to the manufacturer’s guidelines. Quality control testing, library preparation, and differential expression analysis were performed by the Spatial Biology and Genomics Core of Children’s Hospital Los Angeles.

Total RNA quality was determined by measuring RNA Integrity Number using an Agilent Bioanalyzer RNA Pico chip. Transcriptome libraries were prepared using the NEBNext Ultra II Directional RNA Library Prep Kit with single indexing according to the manufacturer’s protocol. Library quality was determined using an Agilent Bioanalyzer DNA 1000 chip. Libraries were sequenced using paired-end 150 bp × 150 bp chemistry on the Illumina HiSeq platform.

Reads were mapped to the UCSC mm10 transcript set using Bowtie2 version 2.1.0, and gene counts were estimated using RSEM v1.2.15. The TMM method in the edgeR R package was used for gene-count normalization, and differentially expressed genes (DEGs) were identified using edgeR. Genes showing altered expression with p < 0.05 and greater than 1.5-fold change were considered differentially expressed. An additional Wilcoxon rank-sum test was applied to reduce effects driven by outliers and focus subsequent analyses on the highest-confidence DEGs.

Raw and processed RNA-seq data generated in this study have been deposited in the Gene Expression Omnibus (GEO) under accession number GSE320302.

### Developmental trajectory analysis

To examine the developmental trajectory of neurodevelopmental processes occurring within the nodose ganglion over time, raw sequencing data from P21 and P35 samples generated in the current study were combined with control nodose ganglion samples collected by our lab in previous studies at P7, P50, and P90. Previously generated datasets were from standard chow-reared pups with maternal care as usual. The P7 and adult datasets were generated previously and are described in McCoy and Kamitakahara^14^.

Using the DESeq2 package in R, principal component analysis was used to examine relationships between groups across the ages assayed. Because P7, P21, and P90 were the ages exhibiting the greatest distance from each other in principal-component space, these ages were used to further examine how gene expression related to specific developmental processes changes over time.

Developmental processes were assessed by comparing gene expression within the following key domains: Cell Cycle/Proliferation, Cell Death/Apoptosis, Myelin, Extracellular Matrix, Cytoskeletal Organization, Ion Pumps and Ion Channels, Synaptic Vesicles, Postsynaptic Density and Ligand Gated Ion Channels, G Proteins, and Cytokine Receptors. Gene lists were curated as an index of gene expression related to each process within each key domain. The curated lists were comparatively conservative and focused on canonical genes directly involved in each process. For example, the synaptic vesicle list consisted of genes encoding synaptophysin, synaptotagmins, synapsins, syntaxins, vesicle-associated membrane proteins, and Snap25. Genes involved in the negative regulation of a process were avoided. Specific genes in each domain are available in Supporting Document 1.

Within each key domain, each gene was classified as significantly increasing, significantly decreasing, or showing no change between P7 and P21 and between P21 and P90 based on FDR <0.05 and an absolute fold change >1.2. Combining the differential expression data between these pairwise age comparisons yielded eight potential outcomes: no change, continuous increase, continuous decrease, peak, valley, early increase, early decrease, late increase, and late decrease. These patterns describe how gene expression within each domain changes over time. Measurements included the proportion of genes within each domain following each pattern, the proportion of transcripts within each domain following each pattern, and the summed expression of transcripts within each domain at each age. One-way ANOVA was used to determine whether summed expression differed between P7, P21, and P90. Additional comparisons at P21 were made between control and HFD^EARLY^ mice using unpaired t-tests.

### Dual in situ hybridization and immunohistochemistry

Right nodose ganglia were collected from male P35 mice during tissue collection following deep anesthesia with ketamine/xylazine (100:10 mg/kg, i.p.) and transcardial perfusion with 0.9% saline followed by 4% paraformaldehyde in 0.1 M PBS, pH 7.4. Tissues were post-fixed for 2 h, cryoprotected overnight in 20% sucrose in PBS, embedded in OCT, frozen, and sectioned at 20 µm. Nodose ganglia were collected in five series on SuperFrost Plus slides for *in situ* hybridization.

Slides were processed using the RNAscope HiPlex v2 assay from Bio-Techne (catalog #324443) with Protease Plus pretreatment. Slides were processed for HiPlex probe detection using the following 12 mouse probes: T1-*Trpa1*, T2-*Cckar*, T3-*Vip*, T4-*Npy2r*, T5-*Lepr*, T6-*Oxtr*, T7-*Sst*, T8-*Htr3b*, T9-*Glp1r*, T10-*Gpr65*, T11-*Calca*, and T12-*Cartpt*. Probes were detected in four sequential rounds (T1-T3, T4-T6, T7-T9, and T10-T12). A DAPI counterstain was included during the imaging rounds and used as an anchor for image registration.

Following detection of all probes, slides were processed for immunohistochemistry to detect the boundaries of neuronal cell bodies. Slides were washed twice for 5 min in ultrapure distilled water, followed by washes in PBS and PBS containing 0.1% Tween-20 (PBST), and incubated for 30 min in a blocking buffer containing 10% normal donkey serum (Jackson ImmunoResearch) and 1% bovine serum albumin (BSA) in PBS. Slides were incubated overnight at room temperature in antibody solution containing 1% BSA in PBS with primary mouse anti-HuC/D antibody (catalog #A-21271, Thermo Fisher Scientific; RRID: AB_221448). Following two washes in PBST, slides were incubated for 1 h at room temperature in antibody solution containing 1% BSA in PBS with secondary Alexa Fluor 594 AffiniPure F(ab’)2 Fragment Donkey Anti-Mouse IgG (Jackson ImmunoResearch, catalog #715586-151; RRID: AB_2340858). Slides were washed with PBST, counterstained with DAPI, coverslipped with ProLong Gold Antifade Mountant (catalog #P36930, Thermo Fisher Scientific), and imaged by confocal microscopy.

For separate synaptophysin immunohistochemistry experiments, brainstem tissue was collected from P21 mice using the same anesthesia, transcardial perfusion, postfixation, cryoprotection, OCT embedding, and 20-µm sectioning procedure described above. Brainstem sections were collected in five series. Intermediate NTS sections were used for synaptophysin quantification.

Slides were washed three times for 5 min in PBS and incubated for 1 h in blocking buffer containing 10% normal donkey serum (Jackson ImmunoResearch), 1% BSA, and 0.75% Triton X-100 in PBS. Slides were incubated overnight at room temperature in antibody solution containing 1% BSA in PBS with primary Synaptophysin (7H12) Mouse Monoclonal Antibody (catalog #9020, Cell Signaling Technology) diluted 1:500. Following five washes for 5 min in PBS, slides were incubated for 1.5 h at room temperature in antibody solution containing 1% BSA in PBS with secondary Alexa Fluor 647 AffiniPure Donkey Anti-Mouse IgG (H+L) (Jackson ImmunoResearch, catalog #715-605-151; RRID: AB_2340863) diluted 1:500. Slides were washed twice for 5 min in PBS, counterstained with NeuroTrace 435/455 Blue Fluorescent Nissl Stain (catalog #N21479, Invitrogen/Thermo Fisher Scientific), and washed four times for 5 min in PBS. Slides were coverslipped with ProLong Gold Antifade Mountant (catalog #P36930, Thermo Fisher Scientific).

### Image acquisition

All images were captured on a Leica STELLARIS laser-scanning confocal microscope using a 40× water-corrected objective. Confocal image stacks were collected through the z-axis at a sampling frequency optimally determined by the Leica LAS X imaging software based on the wavelengths of the fluorophores used for analysis. For nodose ganglia, the two largest sections through the center of the ganglion, separated by 100 µm, were selected for imaging in each sample. For synaptophysin analysis, intermediate NTS sections were selected for imaging. Slides were coded so that the operator was blinded to experimental group during imaging and analysis.

### Image analysis

Confocal image stacks collected following dual *in situ* hybridization and immunohistochemistry were analyzed using the Single-Cell Automated Multiplex Pipeline for RNA (SCAMPR)^76^ to quantify and spatially map highly multiplexed mRNA labeling at single-neuron resolution. Image stacks corresponding to each tissue section were projected across the z-axis and aligned using ImageJ’s *Register Virtual Stack Slices* and *Transform Virtual Stack Slices* plugins, with DAPI serving as the common signal for alignment across rounds. The Cellpose algorithm was used in Google Colaboratory to generate cellular regions of interest (ROIs) based on HuC/D labeling of neuronal cell bodies. Cellular ROIs were imported into ImageJ and manually corrected for inappropriately assigned cell profiles. Corrected ROIs and registered mRNA images were then analyzed using SCAMPR’s semi-automated thresholding macro, which determines the ideal threshold for each image based on a user-defined threshold for a single representative image, maximizing mRNA signal and removing noise. The macro binarized the images, quantified the area fraction of mRNA signal inside each cellular ROI, and exported the results as a gene-by-cell matrix for further statistical analysis in R and GraphPad Prism.

For synaptophysin analysis, images were opened in ImageJ and an ROI was manually drawn around the intermediate NTS using NeuroTrace staining to identify anatomical boundaries. The area fraction of synaptophysin-immunoreactive puncta within the NTS ROI was quantified. Analysts were blinded to experimental group during analysis.

## QUANTIFICATION AND STATISTICAL ANALYSIS

Statistical analyses were performed using R and GraphPad Prism. Unless otherwise indicated, n represents individual animals. Data are presented as mean ± SEM. Statistical significance was defined as p < 0.05. Exact sample sizes, the meaning of n, and the statistical tests used for individual comparisons are reported in the corresponding figure legends.

For the developmental CCK-satiation and glucose preference analyses, one-way ANOVA with Dunnett’s correction was used for comparisons of developmental ages with the mature P120 response, with Tukey correction used for the corresponding glucose preference analysis where indicated. For dietary group comparisons and body composition measurements, two-tailed unpaired t-tests and two-way mixed-effects ANOVA were used as appropriate. For the calorie-dense liquid diet challenge and nutrient preference experiments involving repeated intake measurements, two-way mixed-effects ANOVA was used. For licking microstructure analyses, repeated-measures two-way ANOVA with post hoc Tukey correction was used to assess licking rate across time, and two-tailed unpaired t-tests were used for cumulative licks, burst number, and licks per burst. For brief access taste tests, repeated-measures two-way ANOVA with post hoc Tukey correction was used to assess lick scores across tastant concentrations. For HiPlex RNAscope analyses, two-tailed unpaired t-tests were used to compare mean expression and the proportion of neurons expressing each gene between groups.

For RNA-seq analyses, the TMM method in edgeR was used for gene-count normalization, and edgeR was used to identify differentially expressed genes. Genes with p < 0.05 and >1.5-fold change were considered differentially expressed. A Wilcoxon rank-sum test was additionally applied to reduce effects driven by outliers. For developmental trajectory analyses, DESeq2 was used for principal component analysis and differential expression analysis. Genes were classified as significantly increasing or decreasing between developmental stages using FDR <0.05 and an absolute fold change >1.2. One-way ANOVA was used to compare summed expression across P7, P21, and P90, and unpaired t-tests were used for comparisons between control and HFDEARLY mice at P21.

For synaptophysin analysis, area fraction of synaptophysin-immunoreactive puncta within the intermediate NTS was quantified in 5-7 animals per group and compared using a two-tailed unpaired t-test.

Individual inclusion and exclusion criteria are described in the relevant Method Details sections, including the training criterion for nutrient acceptance testing and the minimum number of trials required for brief access taste-test analyses.

## Supplemental Figures

**Supplemental Figure 1.**
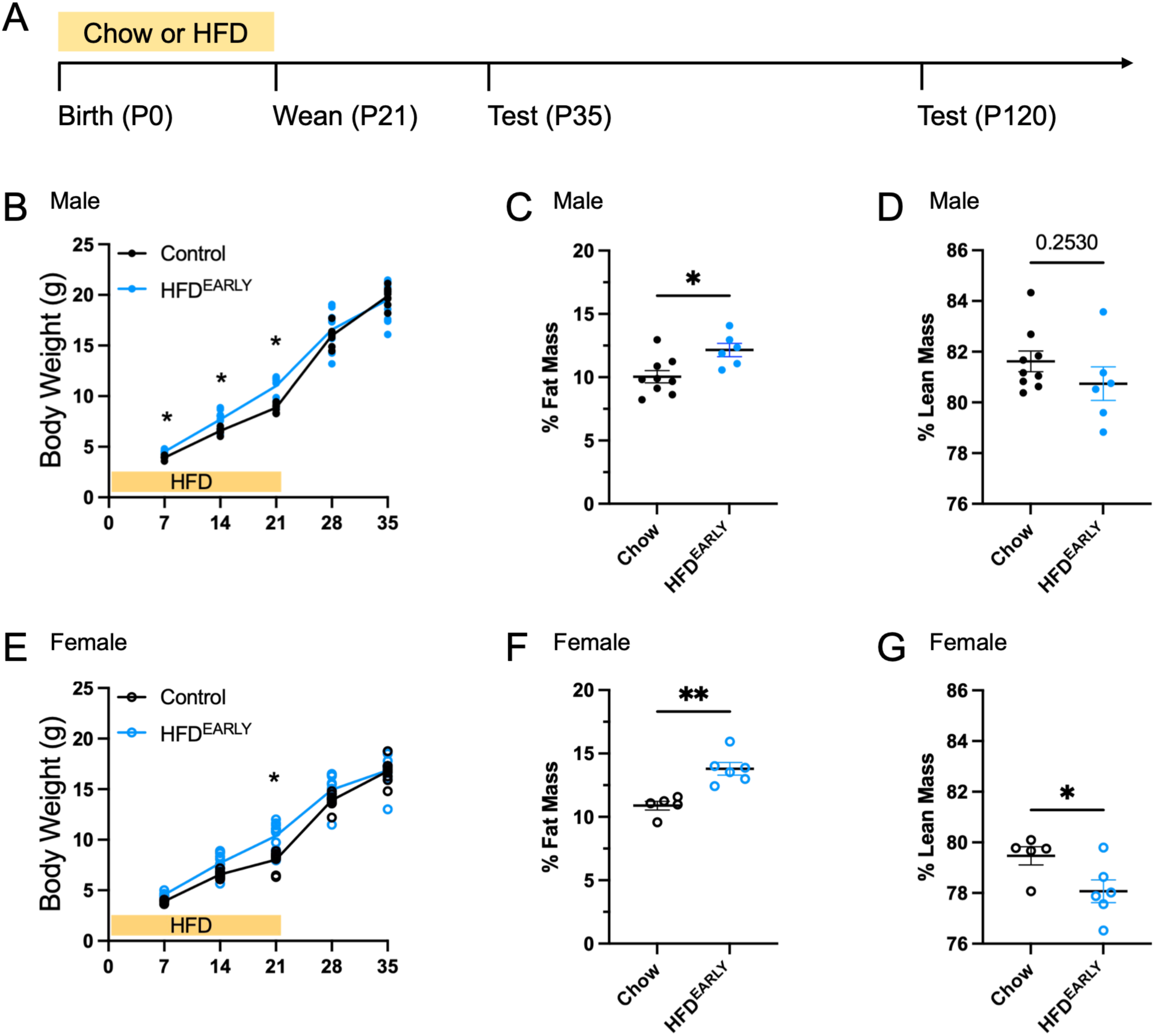
Early life HFD model and effects on body weight and body composition. A) Timeline for dietary manipulation and testing. B) Body weight gain through P35 in male control and HFD^EARLY^ mice. C) Male fat mass as a percentage of total body weight at P35. D) Male lean mass as a percentage of total body weight at P35. E) Body weight gain through P35 in female control and HFD^EARLY^ mice. F) Female fat mass as a percentage of total body weight at P35. G) Female lean mass as a percentage of total body weight at P35. Each point in body weight graphs is the mean value for each sex within a single litter (n=5-11 litters). A single mouse from each litter was assessed for body composition (n=5-9 mice). A two-way mixed-effects ANOVA was used to analyze differences in body weight between groups over time (* p<0.05). A two-tailed unpaired t-test was used to analyze fat and lean mass differences between groups (* p<0.05, ** p<0.01).

**Supplemental Figure 2.**
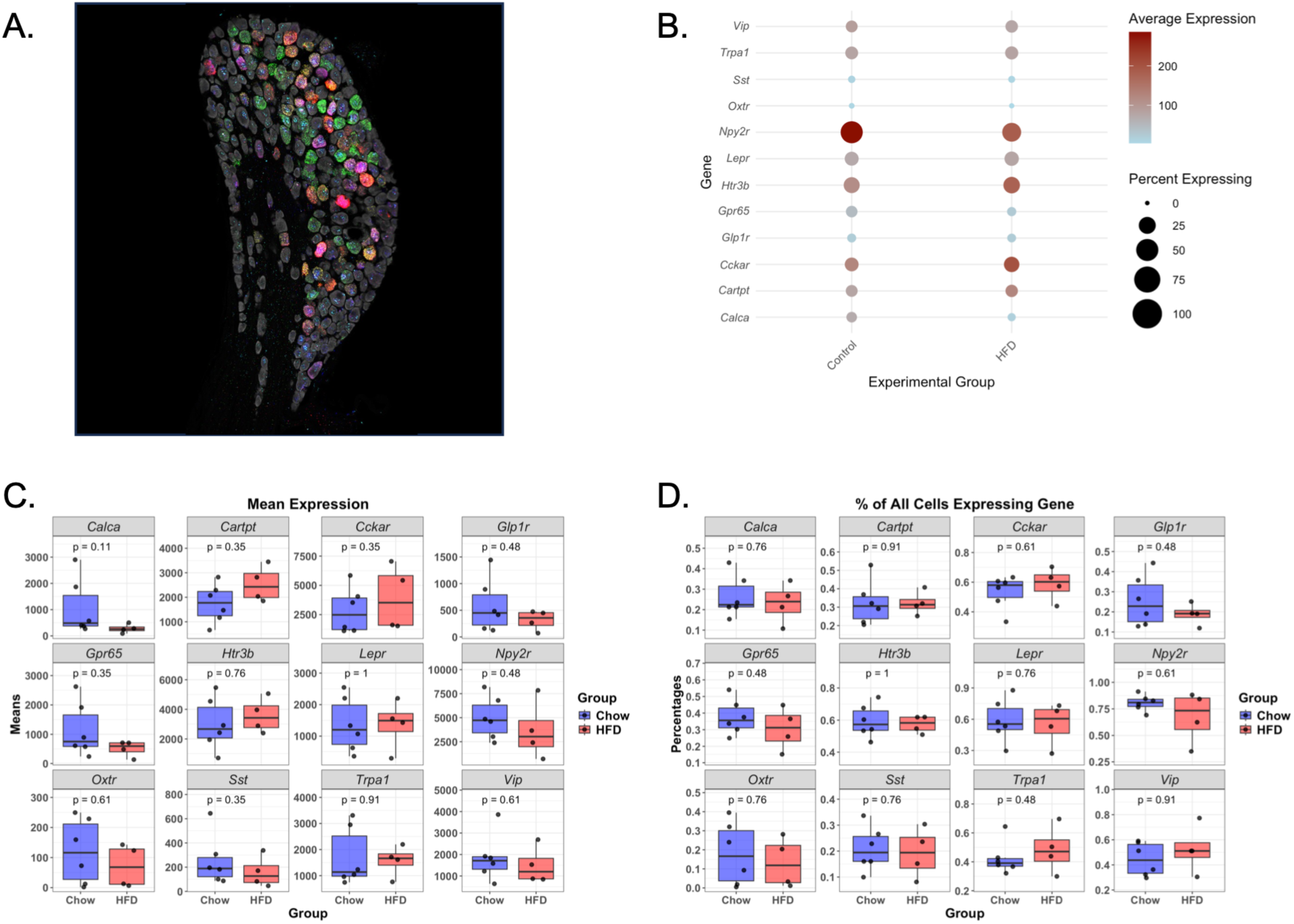
Highly multiplexed in situ hybridization for CCK receptor and other neuronal cell type markers. A) Representative image of probe labeling in a P35 nodose ganglion following image registration. B) Dot plot showing average expression and percentage of cells expressing each gene in control (Chow) and HFD^EARLY^ (HFD) mice. C) Mean expression per neuron for each gene in control and HFD^EARLY^ mice. D) Proportion of neurons expressing each gene in control and HFD^EARLY^ mice. (n=4-6 per group.) A two-tailed unpaired t-test was used to analyze differences in mean expression and percent expressing between groups.

**Supplemental Figure 3.**
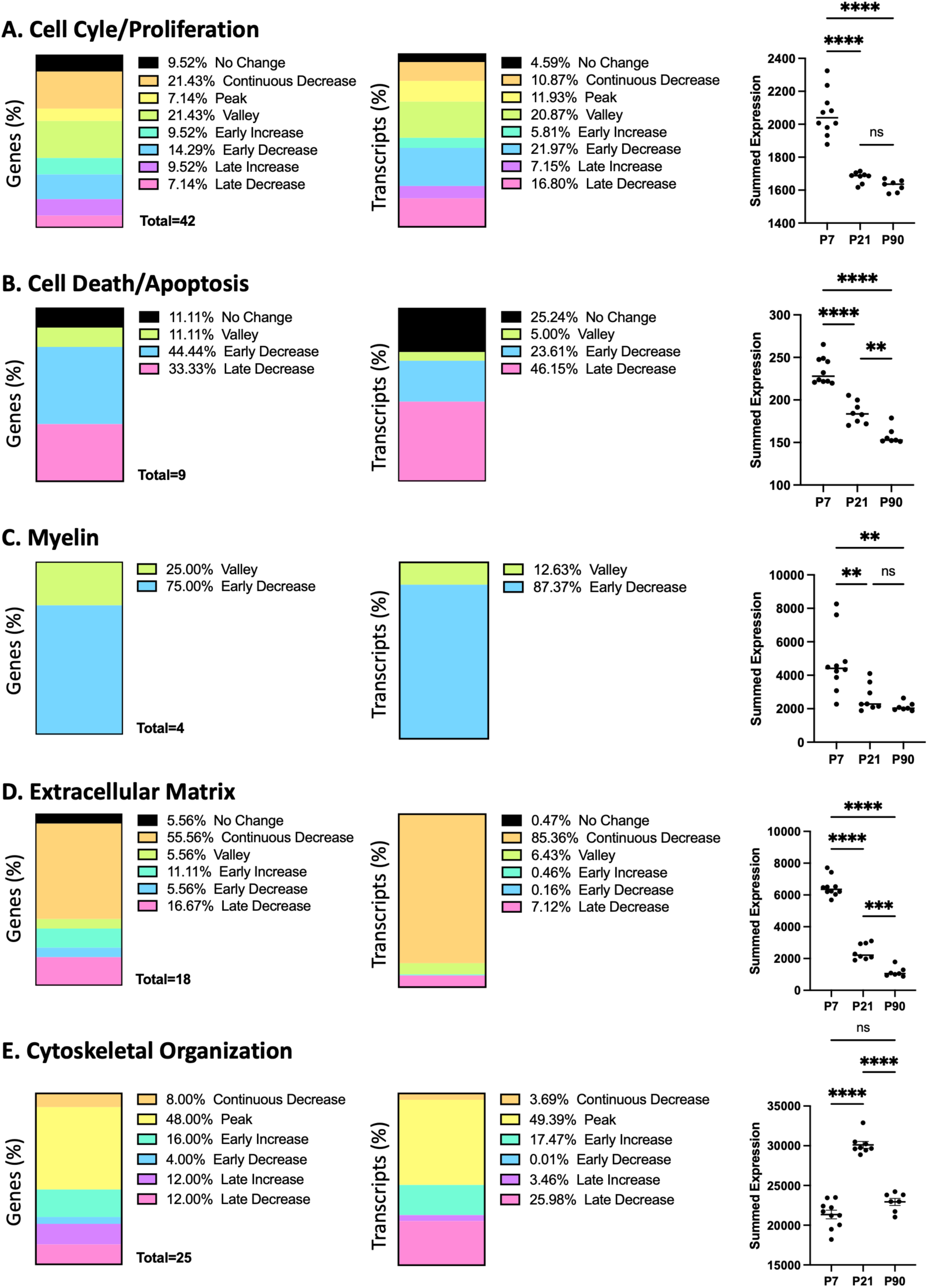
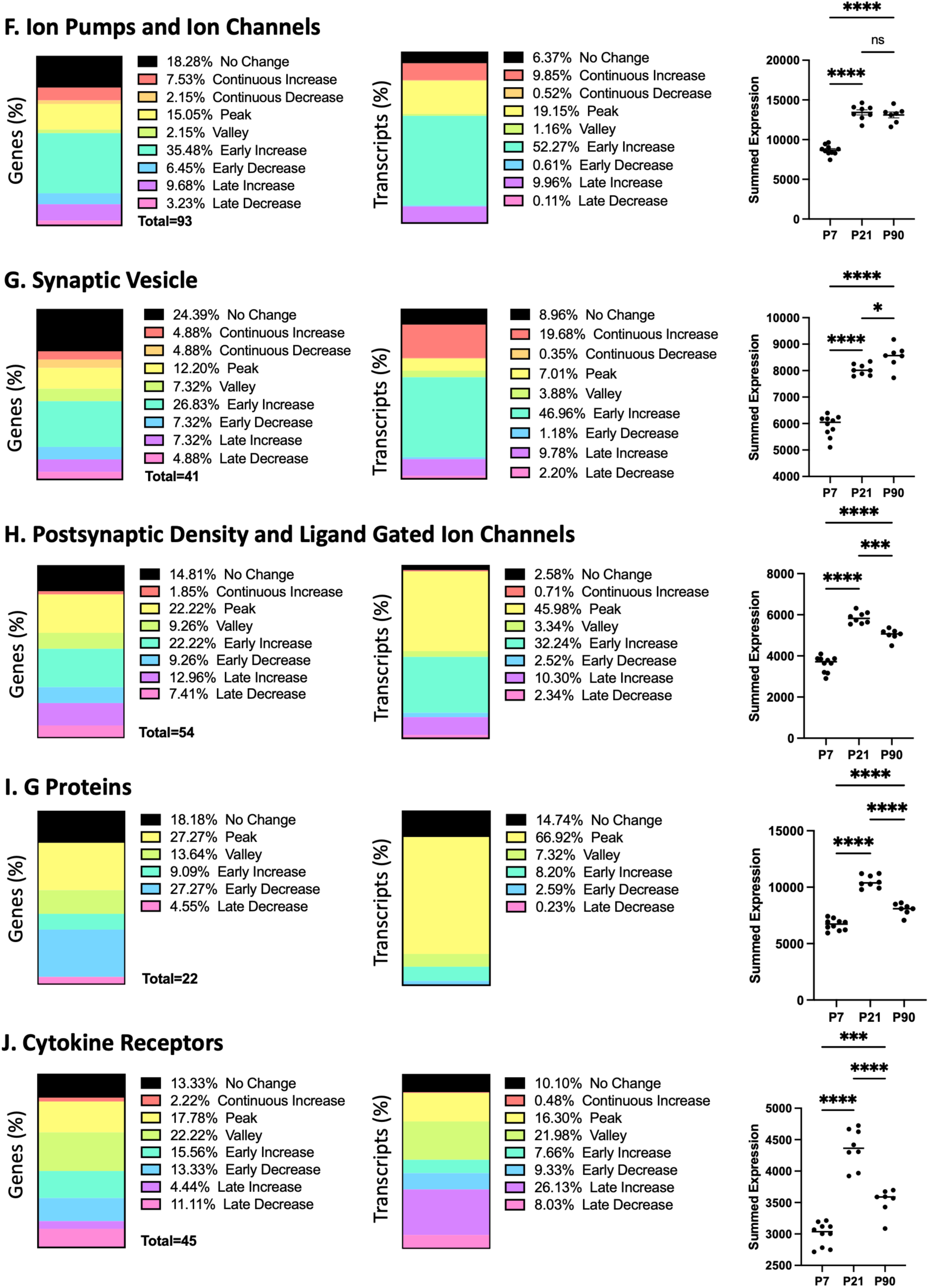
Developmental analysis of nodose ganglion gene expression. Developmental processes were assessed by comparing gene expression within the following key domains: A) Cell Cycle/Proliferation, B) Cell Death/Apoptosis, C) Myelin, D) Extracellular Matrix, E) Cytoskeletal Organization, F) Ion Pumps and Ion Channels, G) Synaptic Vesicles, H) Postsynaptic Density and Ligand-Gated Ion Channels, I) G Proteins, and J) Cytokine Receptors. Stacked bar charts represent the fraction of genes (left column) or transcripts (middle column) exhibiting each expression pattern between P7, P21, and P90. Combining the differential expression data from pairwise comparisons (P7 vs. P21 and P21 vs. P90) yields 8 potential outcomes: no change, continuous increase, continuous decrease, peak, valley, early increase, early decrease, late increase, and late decrease. These patterns describe how gene expression within each domain changes over time. In addition, the summed expression of transcripts within each domain at each age is shown in the right column (n=7-10). A one-way ANOVA was used to determine whether summed expression differed between P7, P21, and P90 (* p<0.05; ** p<0.01; *** p<0.001; **** p<0.0001; ns, not significant).

**Supplemental Figure 4.**
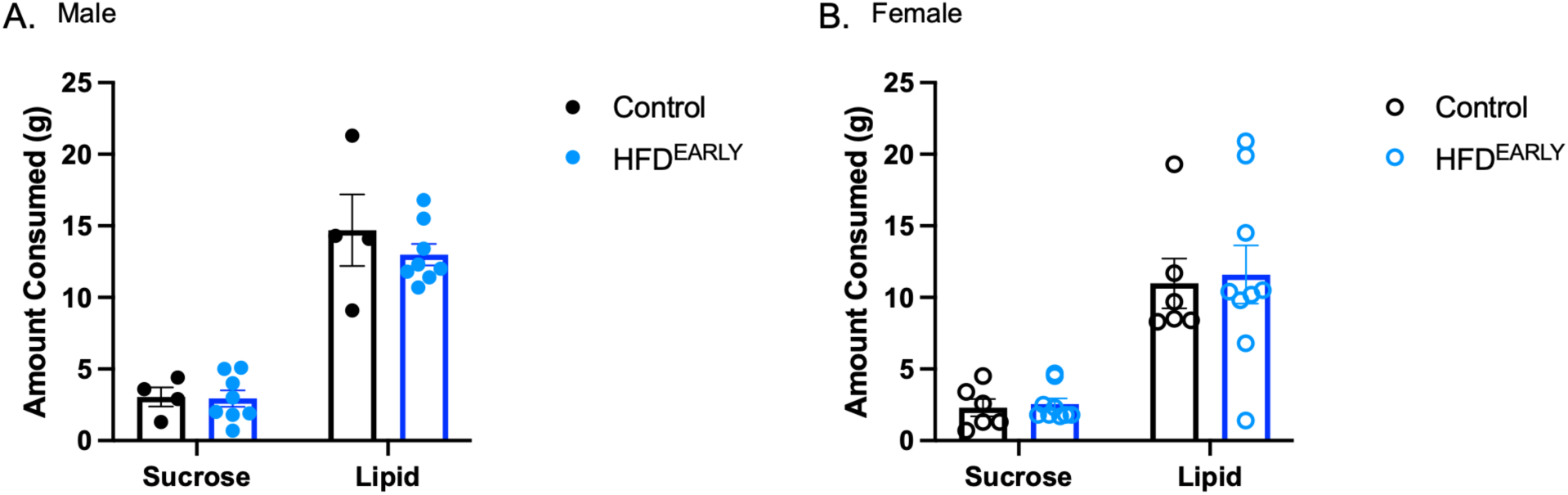
Early life lipid and sucrose consumption does not differ between groups at P35. Two-bottle preference testing (0.6% Intralipid vs. 1.5% sucrose, calorie-matched at 0.06 kcal/mL) was conducted at P35 in male (A) and female (B) control and HFD^EARLY^ mice. No significant differences were observed in the amount of sucrose or lipid consumed between groups in either sex. A two-way mixed-effects ANOVA was used to analyze differences in the amount consumed between groups. ns, not significant.

